# Separable downmodulation of meiotic axis protein deposition and DNA break induction at chromosome ends

**DOI:** 10.1101/2025.02.27.640173

**Authors:** Adhithi R. Raghavan, Kieron May, Vijayalakshmi V. Subramanian, Hannah G. Blitzblau, Neem J. Patel, Jonathan Houseley, Andreas Hochwagen

## Abstract

In many organisms, meiotic crossover recombination is suppressed near the extreme ends of chromosomes. Here we show that multiple, often chromosome-specific, suppressive mechanisms with differing ranges contribute to the consistently low enrichment of recombination-promoting axis proteins and downregulation of DNA double-strand breaks (DSBs) within 20 kb of telomeres in *Saccharomyces cerevisiae*. Suppression of axis proteins is associated with *cis*-encoded signals and correlates with reduced coding density, although whether this sequence feature actively drives suppression remains to be determined. In addition, axis protein suppression requires the histone methyltransferase Dot1 and the Sir silencing complex. We show that Dot1 suppresses Sir complex activity at least in part independently of its canonical target, H3K79, to downmodulate axis protein deposition near chromosome ends. In parallel, the Sir complex, but not Dot1, suppresses the induction of DSBs at a small number of cryptic hotspots by limiting the openness of promoters, the preferred sites of meiotic DSB formation. Much of the reduced DSB induction near chromosome ends persists in *dot1* and *sir3* mutants, indicating that additional layers of regulation contribute to the robust reduction of meiotic recombination effectors near chromosome ends.

## Introduction

Meiosis is a specialized type of cell division that generates haploid gametes from diploid progenitor cells and plays an essential role in promoting genetic diversity ^1^. Meiosis begins with a single round of DNA replication followed by two rounds of chromosome segregation, first segregating homologous chromosomes and then sister chromatids ^2,3^. Accurate segregation during meiosis I relies on meiotic crossover recombination, which exchanges DNA between homologous chromosome pairs and, together with sister chromatid cohesion, forms a physical connection between them ^4^. Meiotic recombination is initiated by programmed DNA double-strand breaks (DSBs), which are catalyzed by Spo11, a highly conserved topoisomerase-like enzyme ^5^.

Although DSBs can occur throughout the genome, their frequencies vary widely. Chromosomal regions classified as “hot” experience frequent DSBs, whereas “cold” regions, including centromeres and chromosome ends, rarely undergo breakage ^6^. Hotness results from a complex interplay of factors, including local base composition, chromatin modifications, meiotic chromosome architecture, as well as environmental factors ^5,7–9^.

In addition to DSBs, the differential distribution of meiosis-specific axis proteins, which localize at the base of the loop-axis structure of meiotic chromosomes, has a major impact on recombination levels across the genome. Axis proteins can promote DSB activity and are also essential for directing repair toward the homologous chromosome, and thus crossover formation ^6,10–12^. In *S. cerevisiae*, elevated binding of the axis proteins Red1 and Hop1 is often associated with markers of crossover repair ^13,14^. Red1 and Hop1 are recruited to the base of chromatin loops by the meiotic Rec8-cohesin complex but can also bind directly to chromatin through the nucleosome-binding activity of Hop1 ^13,15,16^, leading to two independent modes of axis protein-dependent patterning of meiotic recombination.

In *S. cerevisiae*, levels of DSBs and crossover formation exhibit a distinctive pattern near chromosome ends. Levels are elevated above genome average in the end-adjacent regions (EARs), large chromosomal regions located 20-120 kb from chromosome ends, but notably reduced within ∼20 kb from telomeres ^11,17–20^. The depletion within the last ∼20 kb is likely important for two reasons. First, these regions are enriched for repetitive DNA sequences that are vulnerable to non-allelic homologous recombination (NAHR) and genome rearrangements ^21^. Second, crossovers near chromosome ends are less effective at forming stable connections between homologous chromosomes and may lead to chromosome mis-segregation if they are the only connection between homolog pairs ^22^. Consistent with this notion, recombination events near telomeres are linked to a higher risk of Trisomy 21 (Down syndrome) in humans ^23,24^.

*S. cerevisiae* chromosome ends consist of several distinct genomic domains (**Fig. 1a**) ^25,26^, including the telomerase-templated telomeric repeats that cap chromosome ends and the “telomere-associated sequences”, which contain repetitive Y′ and X elements. In addition, the “subtelomeric domains” extend inward for an average of about 20 kb from telomeres. Subtelomeric domains are relatively gene-poor regions, and genes located in these domains often belong to gene families ^27,28^. In vegetative cells, the telomeric repeats and X elements recruit the Sir2/Sir3/Sir4 histone deacetylase complex, establishing transcriptional silencing. This silent chromatin can spread over limited distances into the subtelomeric domains ^29^. The subtelomeric domains, in turn, are defined by a unique chromatin signature, which includes a relative depletion of active chromatin marks ^28,30^. Whether similar chromatin patterns exist during meiotic recombination is unknown.

**Figure 1:**
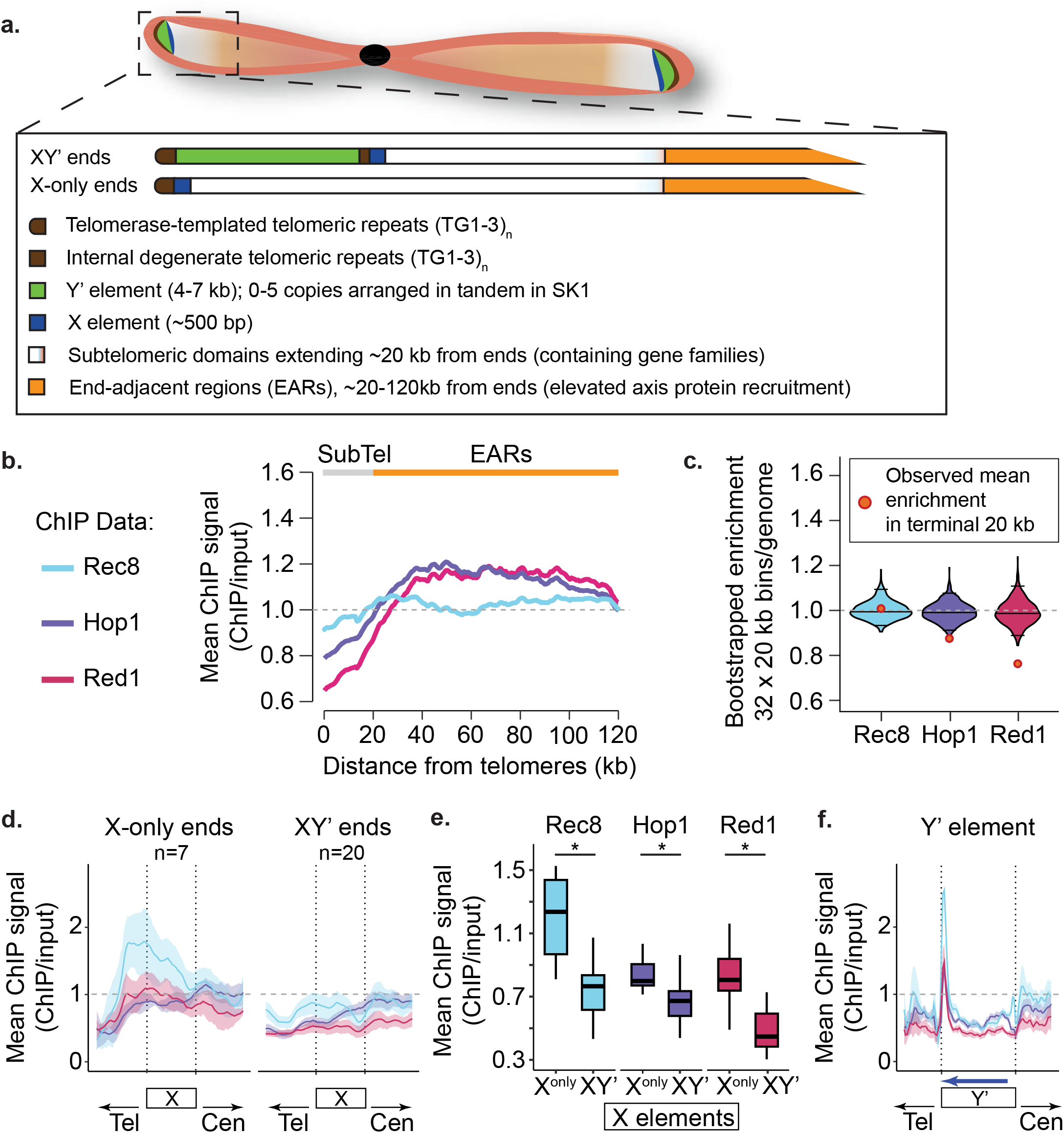
Distinct axis-protein enrichment patterns at chromosome ends. **(a)** Schematic of chromosome-end architecture in *S. cerevisiae*. XY′ ends contain Y′ elements; X-only ends lack Y′ elements. We define the subtelomeric domains as encompassing the last 20 kb from chromosome ends; they thus also encompass any X or Y′ elements. The adjacent EARs extend 20-120kb from ends. **(b)** Mean enrichment of Rec8 (light blue), Red1 (red), and Hop1 (purple) versus distance from telomeres in wild type (WT) early prophase I (T = 3h) from published data ^16,19,36^, normalized to a genome average of 1 (gray dashed line; see Methods: Distance from telomeres plots). Range of the subtelomeric domains (SubTel) is indicated with a solid gray line, EARs are indicated with an orange line. **(c)** Genome-wide bootstrap distributions of fold-enrichment (32 × 20-kb windows; n = 1,000 resamples; see Methods: Bootstrapping plots). Gray dashed line is genome average. Black lines show medians and 95% CIs; orange/red circles mark the observed mean in the last 20 kb. Two-sided empirical test with Benjamini-Hochberg (BH) correction, effect sizes via Cohen’s d (negative = depletion at ends relative to the genome-wide null): Hop1 (p = 0.001; BH = 0.0015; d = −3.51); Red1 (p < 1x10^-6^; BH < 1 x 10^-6^; d = −5.23); Rec8 (p = 0.368; BH = 0.368; d = −0.89). **(d)** Metaplots anchored at X elements, stratified by end class. Only fully annotated X elements were used (X-only, n = 7; XY′, n = 20). Flanks scaled to element length (X: 100% each side). Gray dashed line is genome average, vertical dotted lines mark X boundaries; shaded bands indicate 95% confidence intervals (CI; see Methods: Meta gene analyses, and meta-X and Y’ elements plots). **(e)** Axis protein ChIP signal (Hop1, Red1, Rec8) at X elements on X-only versus XY′ ends. Values represent the mean ChIP/input signal per X element. Box-and-whisker plots show the distribution across elements. Two-sided unpaired Student’s t-tests with BH correction; stars reflect BH-adjusted p (* ≤ 0.05; n.s., not significant). Statistics (per X element; Cohen’s d; positive = higher at X-only): Hop1 (p = 0.0186; BH = 0.0186; d = 1.13); Red1 (p = 0.0058; BH = 0.0118; d = 1.77); Rec8 (p = 0.0079; BH = 0.0118; d = 1.62). **(f)** Metaplot anchored at Y′ elements. Only fully annotated Y′ were analyzed and flanks were scaled to 50% of Y′ length. Blue arrow indicates Y′-ORF orientation. Gray dashed line is genome average and vertical dotted lines mark Y′ boundaries. Shaded bands are 95% CIs (see Methods: Meta gene analyses, and meta-X and Y’ elements plots). Averages of two biological replicates.

Previous studies of meiotic recombination in *S. cerevisiae* have consistently shown a depletion of DSBs in X and Y′ elements as well as in the subtelomeric domains ^11,17,31,32^. While the mechanisms governing DSB suppression in X and Y′ elements remain largely unexplored, the reduced DSB levels in the subtelomeric domains are accompanied by lower recruitment of DSB-promoting factors, such as Rec114 ^33^, and diminished abundance of Hop1 ^19^. Hop1, which recruits Rec114 ^12^, is currently the most upstream regulator of recombination known to be depleted within ∼20 kb from chromosome ends. Whether other axis proteins are similarly depleted and the specific mechanisms controlling this altered distribution, as well as whether depletion is linked to the suppression of meiotic DSBs is unknown.

## Results

### Axis protein enrichment patterns at chromosome ends

Given the reduced axis protein enrichment near chromosome ends and their critical role in meiotic DSB formation and repair, we investigated how axis proteins interact with telomere-associated sequences and subtelomeric domains. These regions are often excluded from sequence-based analyses due to variations in telomere organization even among closely related yeast strains ^25,34,35^, and because the abundance of repetitive sequences and gene families creates challenges for uniquely assigning sequencing reads. Therefore, we tailored our analysis pipeline to account for these unique biological features. To reduce structural mapping artifacts, we mapped reads to high-quality end-to-end chromosome scaffolds of our experimental strain (SK1) ^35^, rather than the commonly used S288c reference. Although the inherent instability of subtelomeric domains means that individual SK1 isolates may differ from the SK1 reference at some ends, this approach eliminates the profound telomere-proximal sequence differences between SK1 and S288c as a major source of error. Further, by exclusively considering optimal matches, we maximized the number of confidently mapped reads, leveraging inherent sequence polymorphisms within telomeric repeat sequences for unique mapping.

To validate our pipeline, we assessed mapping and coverage characteristics in unenriched input datasets. Even in the highly repetitive X and Y′ elements, our pipeline was able to uniquely map 12.8% and 0.87% of single-end reads and 15.2% and 12.8% of paired-end reads, respectively.

In the subtelomeric domains as a whole (within 20 kb from chromosome ends), unique mapping rates were higher, averaging 24.1% for single-end reads and 32.2% for paired-end reads (**Supplementary Fig. 1a**). In addition, we observed no systematic differences in fragmentation efficiency (**Supplementary Fig. 1b**) and only minor differences in sequence coverage due to incomplete X-element annotation (**Supplementary Fig. 1c**), indicating that telomere-proximal regions are sampled comparably to the chromosome interior.

If a read had multiple equally good alignments, our pipeline randomly selected one location as the primary alignment for the read. This approach did not affect metagene analyses but significantly improved signal clarity at individual chromosome ends by closing coverage gaps caused by a lack of polymorphisms. Comparative analysis of ChIP profiles with and without multiple-mapping reads revealed no qualitative differences (**Supplementary Fig. 1d**).

Having validated our pipeline, we analyzed the distribution of axis proteins at chromosome ends using previously published ChIP-seq datasets ^16,19,36^. Meta-analysis showed that Hop1 was depleted below genome average within 20 kb of chromosome ends and enriched above genome average in the EARs (20-120 kb; **Fig. 1b-c**), consistent with previous studies ^19^. This pattern was mirrored by Red1, in line with the observation that Hop1 and Red1 depend on each other for chromosomal association ^16^. Axis protein depletion in the first 20 kb was observed across all chromosome ends and thus could not be attributed to individual telomere outliers (**Supplementary Fig. 2, 3a**).

Yeast chromosome ends are characterized by telomeric repeats, (TG_1–3)n_, and telomere-associated sequences, which include the Y′ and X elements ^37^. The long Y′ elements (4-7 kb) are not found on all ends and can occur as one or multiple copies, whereas a short X element (∼500 bp) is detectable at nearly all chromosome ends ^35,38^ (**Fig. 1a, Supplementary Fig. 2**). Based on these features, we classified chromosome ends into two categories: X-only ends, which contain only the X element, and XY′ ends, which include both X and Y′ elements. Because X elements are relatively small and therefore more difficult to annotate, we classified ends containing a Y′ element as XY′, whereas ends lacking a Y′ element were categorized as X-only. The two ends without annotated X or Y′ elements (**Supplementary Fig. 2**) were included in the X-only category.

Meta-plots of Red1 and Hop1 profiles revealed that axis protein enrichment differed between these two categories. X elements at X-only ends exhibited axis protein enrichment close to the genome average, indicating that isolated X elements have a substantial propensity for axis protein recruitment (**Fig. 1d**). By contrast, X elements at XY′ ends showed Red1 and Hop1 enrichment below the genome average (**Fig. 1d-e**). Meta-analyses revealed an axis protein binding site at the telomere-proximal side of the Y′ element that may act as a local sink for axis protein recruitment (**Fig. 1f**).

The variable presence of Y′ elements also raised the question of whether axis protein depletion is more appropriately assessed by aligning chromosome ends at the X elements. Anchoring on the X elements rather than the telomeres showed a weakened depletion signal in meta-analyses (**Supplementary Fig. 4a**), suggesting that the axis protein depletion in the subtelomeric domains is driven to a substantial degree by sequences located between telomeres and the X elements. Consistent with this interpretation, the mean enrichment of axis proteins in the Y′ elements, which are located in this interval, is significantly below genome average (**Supplementary Fig. 4b**), likely explaining the stronger axis protein depletion on XY′ ends compared to X-only ends (**Supplementary Fig. 4c**).

### Axis protein depletion near telomeres is encoded in *cis*

To directly test whether the local suppression of axis protein binding is due to telomere proximity or intrinsic sequence features, we analyzed chromosome fusions, in which subtelomeric domains were relocated to the chromosome interior ^39^. These chromosomes were constructed in two different genetic backgrounds (S288c and SK1; **Fig. 2a**) and showed overall normal Red1 enrichment, except for regional changes around centromeres, as noted previously ^39^. The need for unique sequences during the fusion process invariably resulted in the deletion of some subtelomeric sequences, but in two cases (S288c: chrI-L and SK1: chrIV-R) the fusion breakpoints were only 6.8 and 11.3 kb from telomeres, allowing for analysis of effects on axis protein enrichment in the subtelomeric domains. At those fused chromosome ends, axis protein distribution mirrored that of native, unfused ends (**Fig. 2b**, **Supplementary Fig. 5**), suggesting that, at least at those ends, reduced axis protein enrichment at chromosome ends is driven by specific cis-encoded sequences rather than by telomere proximity.

**Figure 2:**
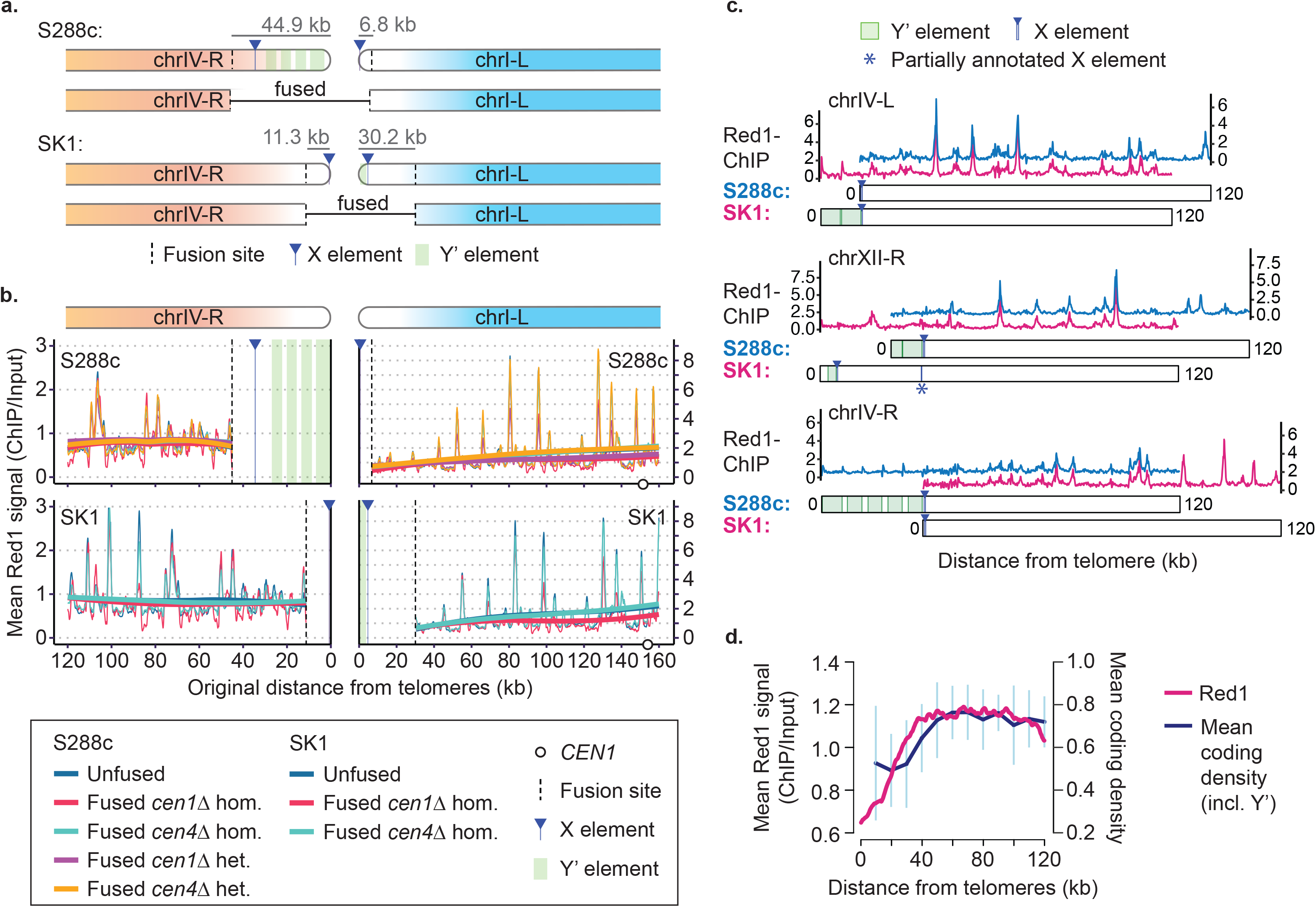
Axis-protein depletion near telomeres is encoded in *cis*. **(a)** Schematic showing the analyzed fusion chromosomes derived either from S288c or SK1 ^39^. The subtelomeric sequences eliminated as part of the fusion process are indicated by grey bars, and the fusion sites, X and Y′ elements are marked. **(b)** Red1 binding (T = 4h) along telomere-proximal arms in SK1/S288c hybrids (chrIV-R and chrI-L), using data from ^39^. Tracks are shown for unfused (WT) and chrIV/I fused strains homozygous (hom.) or heterozygous (het) for the fused chromosomes with either *CEN1* or *CEN4* deleted (colors as indicated). Vertical dashed black line indicates engineered fusion sites. The plot only includes points common to all datasets, which is why the WT unfused data in the regions between the telomeres and the fusion sites are missing. Thin lines are genome-normalized Red1 tracks; thick lines are loess-smoothed overlays for trends (span = 1). Circle indicates *CEN1*. The dip in signal on the right side of chrI-L is because of deletion of *CEN1* in some of the strains as previously described ^39^. (**c**) Example distance-from-telomere plots showing Red1 ChIP/Input signal along 3 matching chromosome arms in WT SK1/S288c hybrid strains, carrying a haploid genome of SK1 and a haploid genome of S288c ^39^. SK1 and S288c sequences are sufficiently different that about 25% of reads can be assigned to one of the two genomes ^36^. Red1 profiles were manually aligned to show that the offset caused by differences in Y′ elements and other subtelomeric sequences does not greatly alter the relative distribution or height of Red1 peaks. See **Supplementary** Fig. 6 for all ends in register. **(d)** Coding density versus distance from telomeres overlaid on a metaplot of Red1 enrichment. Graph points show mean coding density in 10-kb bins, plotted at bin midpoints (right y-axis). Error bars are standard deviation. Line connects the dots for easier visualization of trend. Y′ elements are included and are responsible for the uptick in signal in coding density in the most telomere-proximal bin.

To further test whether reduced axis protein enrichment is encoded in *cis*, we took advantage of the strain-specific differences between the subtelomeres of S288c and SK1 by analyzing Red1 enrichment data from S288c/SK1 hybrids that harbor one haploid chromosome complement from S288c and one complement from SK1 ^39^. Because of the large number of sequence differences between the two genomes, many reads can be uniquely assigned, enabling internally controlled analysis of axis protein enrichment between the two genomes ^36^. Comparison of individual chromosome ends of S288c and SK1 showed that the presence of Y′ elements and other subtelomeric sequences primarily resulted in shifts of Red1 peaks toward the chromosome interior (**Supplementary Figs. 6** **and 7a**). After correcting for this displacement, the relative distribution of Red1 peaks and peak heights remained remarkably similar for matching ends (**Fig. 2c**, **Supplementary Fig. 6** **and 7a**), indicating that changing the distance from the nearest telomere does not affect Red1 binding patterns. However, the distal-most sequences, including Y′ elements and other telomere-associated sequences, were consistently under-enriched for axis proteins (**Supplementary Fig. 7b**), indicating that axis protein enrichment near chromosome ends is to a large extent hardwired into the underlying DNA sequence.

Across the genome, gene-rich regions exhibit overall higher axis protein enrichment ^13,16^. As the subtelomeric domains are comparatively gene-poor ^27,28^, we asked how well local coding density (the fraction of DNA that encodes open reading frames) correlated with axis protein enrichment. This analysis revealed a strong correlation between the regions of reduced coding density and Red1 depletion (**Fig. 2d**). In addition, binning the genome into 20-kb bins showed a positive correlation between coding density and average Red1 binding, similar to the effect seen across the genome (**Supplementary Fig. 4d**). Thus, reduced coding density may partially underlie the reduced axis protein recruitment within 20 kb of telomeres. However, we note that this correlation was only apparent when Y′ elements were excluded from the analysis, perhaps because of unique chromatin signatures on Y′ elements. Interestingly, the correlation was absent in the EARs (**Supplementary Fig. 4d**), presumably reflecting the specialized regulation of axis protein deposition in these regions ^19^.

### Rec8-dependent and independent pathways mediate axis protein localization at chromosome ends

Chromosomal recruitment of Red1 and Hop1 is mediated by two parallel pathways. The meiotic Rec8-cohesin complex preferentially recruits axis proteins to sites of convergent transcription ^16^, whereas the chromatin-binding region (CBR) of Hop1 directs axis proteins to nucleosome-dense regions (“islands”) characterized by higher coding density ^13,16^. We sought to determine whether the depletion of axis proteins near chromosome ends could be attributed to the reduced activity of one of these pathways.

Intriguingly, Rec8-cohesin levels in the subtelomeric domains were comparable to the genome average. Although enrichment trended slightly lower, the depletion was within the 95% confidence interval of a bootstrapped distribution (p = 0.37) and thus was not significantly different from the rest of the genome (**Fig. 1b-c**). Therefore, the depletion of Hop1 and Red1 in the subtelomeric domains is not simply due to reduced Rec8-cohesin binding in these regions.

The regions enriched for Red1 and Hop1 in the telomere-associated sequences were also enriched for Rec8: Rec8 was bound more strongly to the X elements of X-only ends than to XY′ ends (**Fig. 1d-e**) and formed a strong peak at the telomere-proximal side of Y′ elements (**Fig. 1f**). This peak coincided with the 3’ end of the Y′-encoded open reading frame (ORF), consistent with other chromosomal Rec8 peaks, which are typically enriched downstream of ORFs ^16,40^. Previous studies have demonstrated that active transcription can induce the sliding of the cohesin ring and direct axis protein association to the end of ORFs ^16,41,42^. Therefore, transcription of the Y′ ORF may similarly influence axis protein deposition within Y′ elements.

The coincident binding of axis proteins and Rec8 supports the hypothesis that Rec8-cohesin is an important contributor to axis protein deposition at chromosome ends. To directly test this possibility, we examined Red1 binding in a *rec8* mutant strain using spike-in normalized ChIP-seq datasets ^13,36^. In the subtelomeric domains, Red1 levels were significantly reduced in the *rec8* mutant compared to wild type (**Fig. 3a-b**). Importantly, the remaining Red1 enrichment was no longer depleted in the subtelomeric domains. These data suggest that the molecular mechanism suppressing axis protein levels in the subtelomeric domains interferes with Rec8-dependent recruitment.

**Figure 3:**
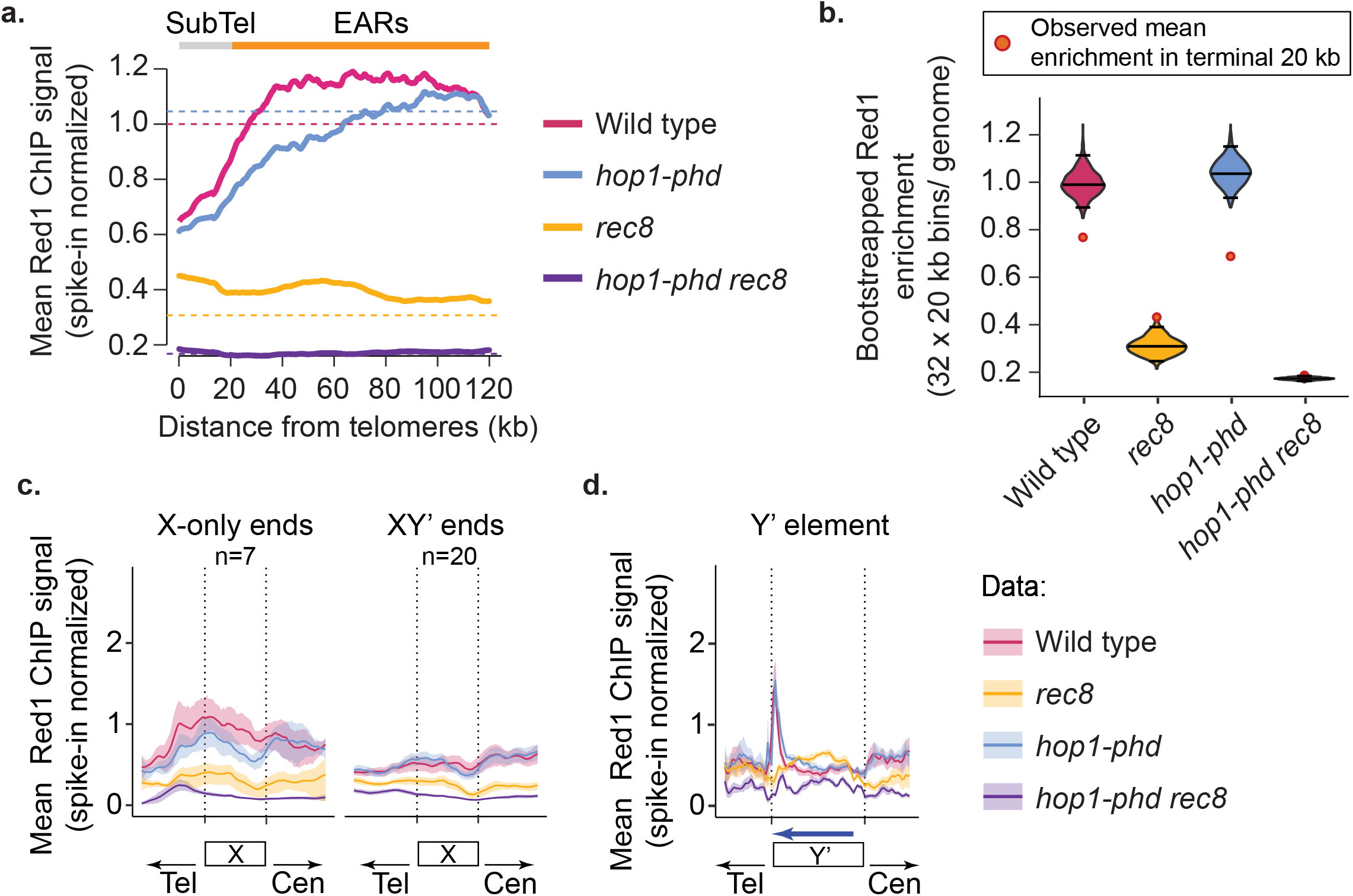
Differential recruitment of Red1 at chromosome ends by Rec8-dependent and Rec8-independent pathways. **(a)** Distance-from-telomere profiles of Red1 (spike-in normalized; see Methods) in WT, *rec8*, *hop1-phd*, and *hop1-phd rec8* during early prophase I (T = 3h) using published data ^13,15^ (see Methods: Distance from telomeres plots). Colored dashed lines indicate each strain’s genome-wide mean after spike-in scaling. Ranges of subtelomeric domains (gray) and EARs (orange) are indicated above the plot. **(b)** Genome-wide bootstrap distributions of fold-enrichment (32 × 20-kb windows; n = 1,000 resamples; see Methods: Bootstrapping plots). Black lines show medians and 95% CIs; orange/red circles mark the observed means in the last 20 kb. Two-sided empirical tests with BH correction; Cohen’s d (negative = depletion at ends): WT (p < 1x10^-6^; BH < 1x10^-6^; d = −5.23); *hop1-phd* (p < 1x10^-6^; BH < 1x10^-6^; d = −7.85); *rec8* (p = 0.001; BH = 0.0013; d = 3.54); *hop1-phd rec8* (p = 0.059; BH = 0.059; d = 1.87). **(c-d)** Meta-X and Y′ elements plots at chromosome ends. X elements were stratified by end class and only fully annotated X elements were used (X-only, n = 7; XY′, n = 20) with flanks scaled to 100% of X length. Y′ elements use flanks scaled to 50% of Y′ length. Blue arrow indicates Y′-ORF orientation. Shaded bands show two-sided 95% CIs (see Methods: Meta gene analyses, and meta-X and Y′ elements plots). Averages of two biological replicates.

On the other hand, analysis of a *hop1-phd* mutant, which lacks the CBR ^13,15^, revealed only minor effects in the subtelomeric domains (**Fig. 3a-b**). However, we consistently noted reduced Red1 binding in the neighboring EARs (**Fig. 3a**), implying a role for the Hop1 CBR in promoting axis protein enrichment in the EARs. In line with the two recruitment mechanisms acting in parallel ^13,15^, the persistent subtelomeric axis protein signal in *rec8* mutants was abolished upon introduction of the *hop1-phd* mutation (**Fig. 3a-b**). Similar additive effects were observed in the X and Y′ sequences (**Fig. 3c-d**). These data indicate that both pathways of axis recruitment contribute additively to Red1 binding at chromosome ends. However, the Rec8-dependent pathway, which is responsible for recruiting the majority of Red1, is the regulatory target for the relative depletion of axis proteins from chromosome ends.

### Dot1 is required for subtelomeric depletion of axis proteins

Subtelomeric domains share qualitative similarities with pericentromeric regions, particularly in the differential enrichment of axis proteins and Rec8-cohesin. Both regions show a relatively higher abundance of Rec8-cohesin compared to Red1, with most data points falling below the genome-wide regression line (**Fig. 4a**). At pericentromeres, this differential enrichment of axis factors persists even if the centromere itself is inactivated ^39^, indicating that the local DNA sequence or chromatin environment influences axis protein binding. Given the distinct chromatin state of the subtelomeric domains, we investigated whether chromatin modifiers specific to these regions contribute to axis protein depletion.

**Figure 4:**
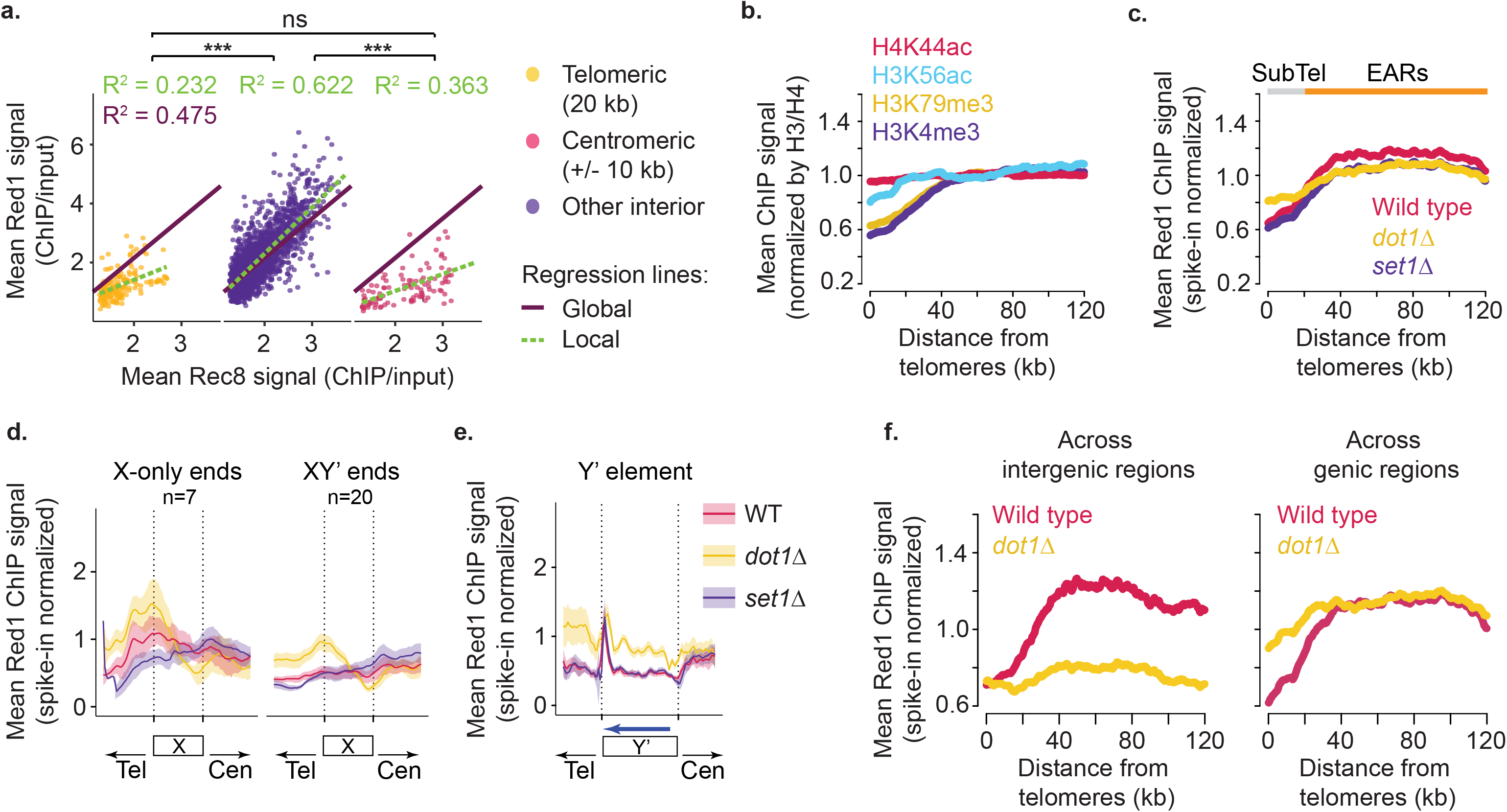
Dot1 shapes axis-protein distribution at chromosome ends and chromosome interiors. **(a)** Mean Red1 vs Rec8 enrichment at Rec8 peaks, split by region (terminal 20 kb, centromeres ±10 kb, interior) using published data ^16^. Global fit (purple) and region-specific fits (green dashes). Slope comparisons (two-sided Student’s t-tests; BH-adjusted): interior vs telomeres (p = 3.43x10^-19^; BH = 5.14x10^-19^); interior vs pericentromeres (p = 3.12x10^-24^; BH = 9.36×10^-24^); telomeres vs pericentromeres (p = 0.525; BH = 0.525). Stars denote BH-adjusted p-values (*** ≤ 0.001; n.s., not significant) (See Methods: Quantification of Red1 and Rec8 signals). **(b)** Mean enrichment of H4K44ac and H3K56ac ^44^, H3K4me3 ^45^, and H3K79me3 versus distance from telomeres, each normalized to H3 or H4 (see Methods: Distance from telomeres plots). **(c)** Spike-in–normalized Red1 distance profiles in WT, *dot1Δ*, and *set1Δ*. Data from ^36^. See Methods: Distance from telomeres plots. Note that *set1Δ* mutants are somewhat less synchronous because of delays in premeiotic DNA replication. **(d-e)** Meta-X and Y′ elements plots of Red1 enrichment in WT (red), *dot1Δ* (yellow) and *set1Δ* mutants (purple). X elements stratified by end class and only fully annotated X elements were used (X-only, n = 7; XY′, n = 20) with flanks scaled to 100% of X length. Y′ elements use flanks scaled to 50% of Y′ length. Blue arrow indicates Y′-ORF orientation. Shaded bands show two-sided 95% CIs (see Methods: Meta gene analyses, and meta-X and Y′ elements plots). **(f)** Region-stratified Red1 enrichment profiles in WT (pink) and *dot1Δ* mutants (yellow) across intergenic (left) and genic (right) sequences versus distance from telomeres. Averages of two biological replicates.

We focused our analysis on histone marks related to meiotic DSB formation or those that are specifically different in the subtelomeric domains ^30,43–46^. ChIP-seq analysis identified two marks, H3K4me3 and H3K79me3, that closely matched the pattern of axis protein depletion (**Fig. 4b**). Both marks are long-lasting indicators of active gene expression that are depleted from subtelomeric domains ^28,30^ and have been implicated in the control of meiotic DSB formation ^43,46^. We note that relative depletion is also apparent when comparing genes in the subtelomeric domains to the rest of the genome by metagene analyses (**Supplementary Fig. 8a**) and thus is not simply due to the lower coding density of the subtelomeric domains.

To explore the role of these histone modifications in axis protein depletion, we used spike-in normalized ChIP-seq ^36^ to analyze Red1 in mutants lacking Set1 or Dot1, the enzymes responsible for the trimethylation of H3K4 and H3K79, respectively. In *set1Δ* mutants, subtelomeric Red1 binding patterns were indistinguishable from wild-type, although we observed an overall reduction in Red1 binding (**Fig. 4c-e, Supplementary Fig. 8b-c**), consistent with previous analyses ^36^. In contrast, *dot1Δ* mutants exhibited a marked redistribution of Red1. Red1 levels were relatively elevated in the Y′ and X elements and no longer significantly different from genome average in the subtelomeric domains (p = 0.16) but were reduced in the EARs (**Fig. 4c-e, Supplementary Fig. 8b-c**), indicating that Dot1 is an important regulator of axis protein distribution. In addition, Red1 levels in *dot1Δ* mutants also trended lower in the pericentromeric regions (p = 0.06 after Benjamini-Hochberg correction; **Supplementary Fig. 8d-e**). This effect is opposite to the increase seen in the subtelomeric domains and indicates that subtelomeric domains and pericentromeres use different mechanisms to achieve a relative depletion of Red1 compared to Rec8-cohesin. Intriguingly, the reduced binding of Red1 in the chromosome interior in *dot1Δ* mutants primarily affected intergenic regions whereas the increase in Red1 association near chromosome ends in *dot1Δ* mutants occurred predominantly on gene bodies (**Fig. 4f**).

To test whether the effects of Dot1 on Red1 recruitment are related to H3K79 trimethylation, we analyzed mutants, in which H3 lysine 79 was changed to arginine (*hht1/2-K79R*) ^46^. These mutants recapitulated the drop in Red1 levels in the EARs (**Fig. 5a**), suggesting that Dot1 promotes axis protein binding in the EARs by methylating H3K79. However, Red1 levels in *hht1/2-K79R* mutants also decreased in the subtelomeric domains and in the Y′ and X elements (**Fig. 5a-d**) and did not decrease in the pericentromeric regions (**Supplementary Fig 8d-e**). These observations reveal an H3K79-independent activity of Dot1 that regulates axis protein enrichment near chromosome ends and centromeres. MNase-seq analysis of *dot1Δ* mutants failed to reveal obvious changes in chromatin accessibility in the subtelomeric domains or the pericentromeres **(Supplementary Fig. 8f-g)**, suggesting that the effects on Red1 recruitment seen in *dot1Δ* mutants are not the result of altered nucleosome occupancy. Together, these data indicate that Dot1 plays multiple roles in controlling axis protein recruitment, only some of which require H3K79.

**Figure 5:**
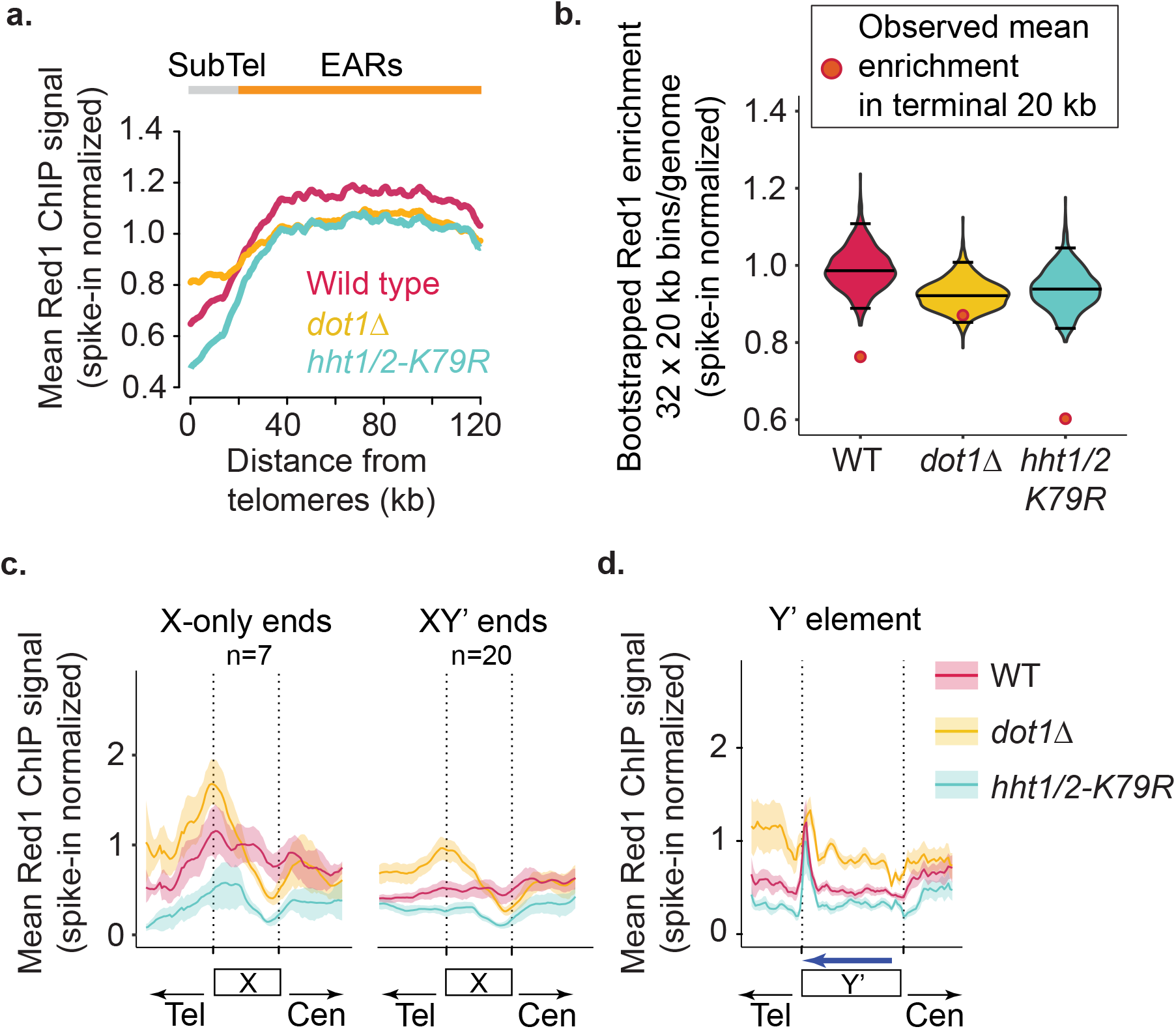
H3K79-independent activity of Dot1 reduces axis proteins near chromosome ends. **(a)** Distance-from-telomere profiles of Red1 enrichment (spike-in normalized) in WT, *dot1Δ,* and *hht1/2-K79R* during early prophase I (T = 3h; see Methods: Distance from telomeres plots). WT and *dot1Δ* data are the same as in Fig. 4c. **(b)** Genome-wide bootstrap distributions (32 × 20-kb windows; n = 1,000 resamples) of Red1 enrichment. Black lines show medians and two-sided 95% CIs; orange/red circles mark the observed mean in the terminal 20 kb. Two-sided empirical test, with BH correction; effect sizes via Cohen’s d: WT (p = 0.0020; BH = 0.0030; d = −4.09); *dot1Δ* (p = 0.180; BH = 0.180; d = −1.30); *hht1/2-K79R* (p = 0.0010; BH = 0.0030; d = −6.24). **(c-d)** Meta-X and Y′ elements plots at chromosome ends. X elements stratified by end class and only fully annotated X elements were used (X-only, n = 7; XY′, n = 20) with flanks scaled to 100% of X length. Y′ elements use flanks scaled to 50% of Y′ length. Blue arrow indicates Y′-ORF orientation. Shaded bands show two-sided 95% CIs (see Methods: Meta gene analyses, and meta-X and Y′ elements plots). Averages of two biological replicates.

### Effects of Dot1 on axis protein deposition depend on Sir3

The Sir complex is an important regulator of telomere-associated chromatin. Indeed, one consequence of losing Dot1 function is a deregulation of the Sir complex ^47,48^. To test whether the Sir complex has a role in telomere-proximal axis protein deposition, we determined the distribution of Red1 in *sir3* mutants using spike-in normalized ChIP-seq analysis. Binding levels of Red1 in the EARs were higher in *sir3* mutants (**Fig. 6a**), consistent with previous analyses of Hop1 in *sir2* mutants ^19^, but meta-analysis revealed no significant difference in the average depletion of Red1 in the last 20 kb between *sir3* and wild-type strains (**Fig. 6a-b**). Similarly, meta-plots of average Red1 profiles on X and Y′ elements in *sir3* mutants showed no new peaks compared to wild-type (**Fig. 6c-d**), although enrichment levels were lower on both types of X element (**Fig. 6c**). Thus, the Sir complex is not a major regulator of telomere-proximal axis protein depletion in the presence of *DOT1*.

**Figure 6:**
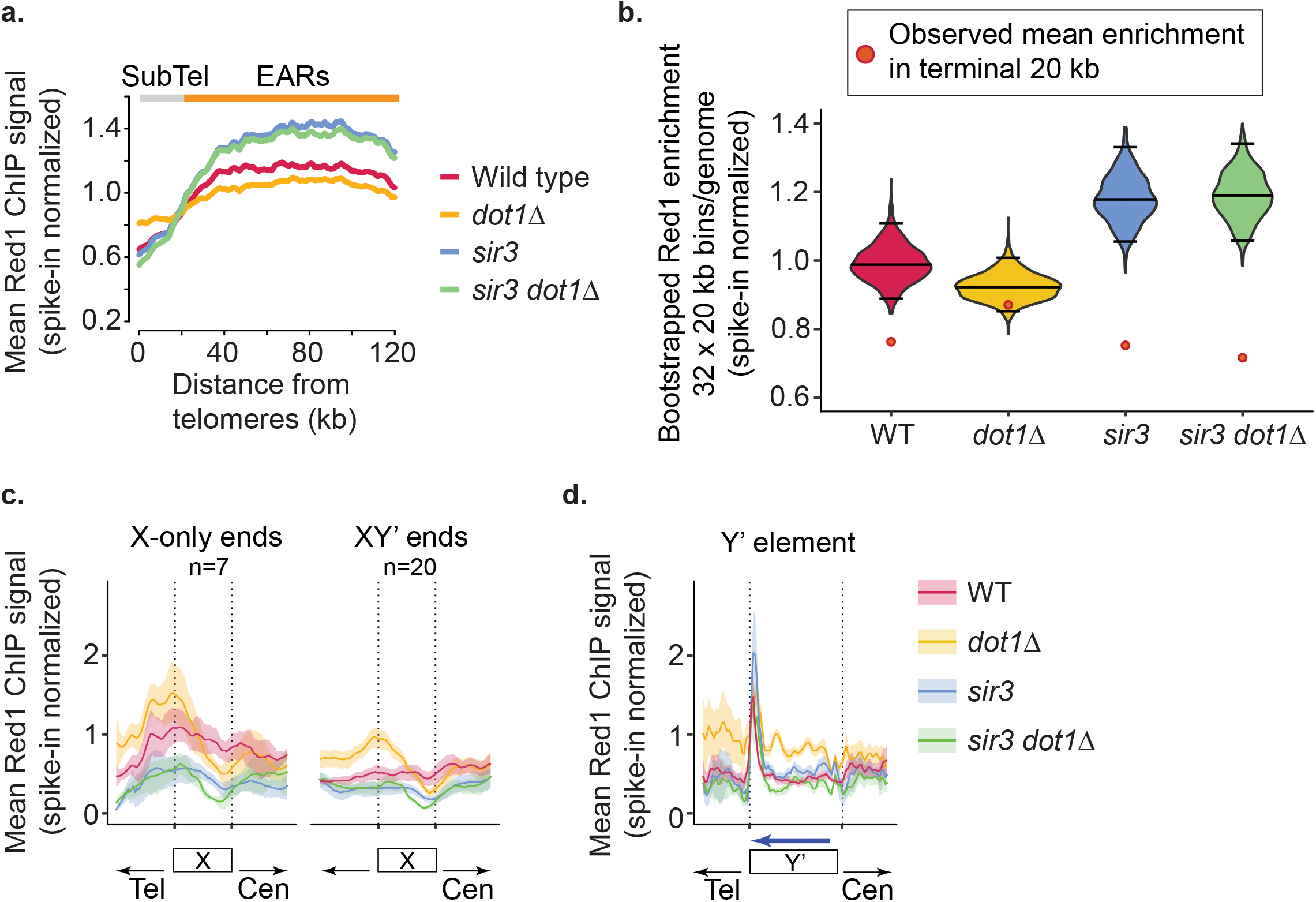
Effects of Dot1 on axis-protein deposition depend on Sir3. **(a)** Distance-from-telomere profiles of Red1 enrichment (spike-in normalized) in WT, *dot1Δ*, *sir3*, and *sir3 dot1Δ* during early prophase I (T = 3h; see Methods: Distance from telomeres plots). **(b)** Genome-wide bootstrap distributions (32 × 20-kb windows; n = 1,000 resamples) of Red1 enrichment. Black lines show medians and two-sided 95% CIs; orange/red circles mark the observed mean in the terminal 20 kb. Two-sided empirical test, with BH correction; effect sizes via Cohen’s d: WT (p = 0.002; BH = 0.003; d = −4.09); *dot1Δ* (p = 0.18; BH = 0.18; d = −1.30); *sir3* (p = 0.001; BH = 0.002; d = −5.87); *sir3 dot1Δ* (p = 0.001; BH = 0.002; d = −6.50). **(c-d)** Meta-X and Y′ elements plots at chromosome ends. X elements stratified by end class and only fully annotated X elements were used (X-only, n = 7; XY′, n = 20) with flanks scaled to 100% of X length. Y′ elements use flanks scaled to 50% of Y′ length. Blue arrow indicates Y′-ORF orientation. Shaded bands show two-sided 95% CIs (see Methods: Metagene analyses, and meta-X and Y′ elements plots). Averages of two biological replicates.

Intriguingly, however, mutation of *sir3* reversed many of the Red1 recruitment phenotypes of *dot1Δ* mutants, including the increased Red1 binding in the subtelomeric domains and the decreased binding in the EARs and pericentromeres (**Fig. 6a-b, Supplementary Fig. 9a-b**). Moreover, the binding pattern of Red1 across X and Y′ elements in the *sir3 dot1Δ* double mutant matched the phenotype of the *sir3* mutant rather than the *dot1Δ* mutant (**Fig. 6a-b**). This phenotypic epistasis suggests that Dot1 either counteracts the activity of the Sir complex to control Red1 distribution or else that absence of *DOT1* creates an environment in which the Sir complex can influence axis protein recruitment.

### Conservation of Sir-dependent telomeric heterochromatin during meiotic recombination

To better understand the role of the Sir complex in meiosis, we analyzed the genomic distribution of Sir3 in wild-type cells by ChIP-seq. In vegetative cells, the Sir complex establishes silent chromatin domains at mating-type loci and telomere-associated sequences ^49,50^. At some chromosome ends, Sir chromatin also spreads from the X element into the neighboring subtelomeric domains ^29,51^, although spreading across entire subtelomeric domains is only observed under conditions of Sir overexpression ^30^.

Our analyses at the time of meiotic induction and during the peak of DSB formation (3 hours post-induction) revealed consistent Sir3 binding patterns (**Fig. 7a, Supplementary Fig. 10**). In addition to the telomere-proximal silent mating type loci on chrIII (*HML*, *HMR*), Sir3 was enriched at both types of X elements, although binding on X elements at XY′ ends was lower than that observed at X-only ends (**Fig. 7a-b**). Sir3 was also enriched upstream of the 5’ end of the Y′-encoded ORF (**Fig. 7c**). In addition, we observed limited and heterogeneous spreading of Sir3 from subtelomeric X elements into adjacent domains on certain chromosome ends. For example, on chrVII-L, Sir3 spread approximately 6 kb from the X element during early prophase I (3 hours post-induction), whereas no detectable spreading occurred on chrVI-L (**Fig. 7d-e**). Spreading distances varied across chromosome ends (**Supplementary Fig. 11a-b**) and did not correlate with the presence of Y′ elements (**Supplementary Fig. 10**), mirroring the behavior of Sir proteins in vegetative cells ^29,30^. These findings indicate that Sir-dependent telomeric heterochromatin remains largely unchanged as cells initiate meiotic recombination.

**Figure 7:**
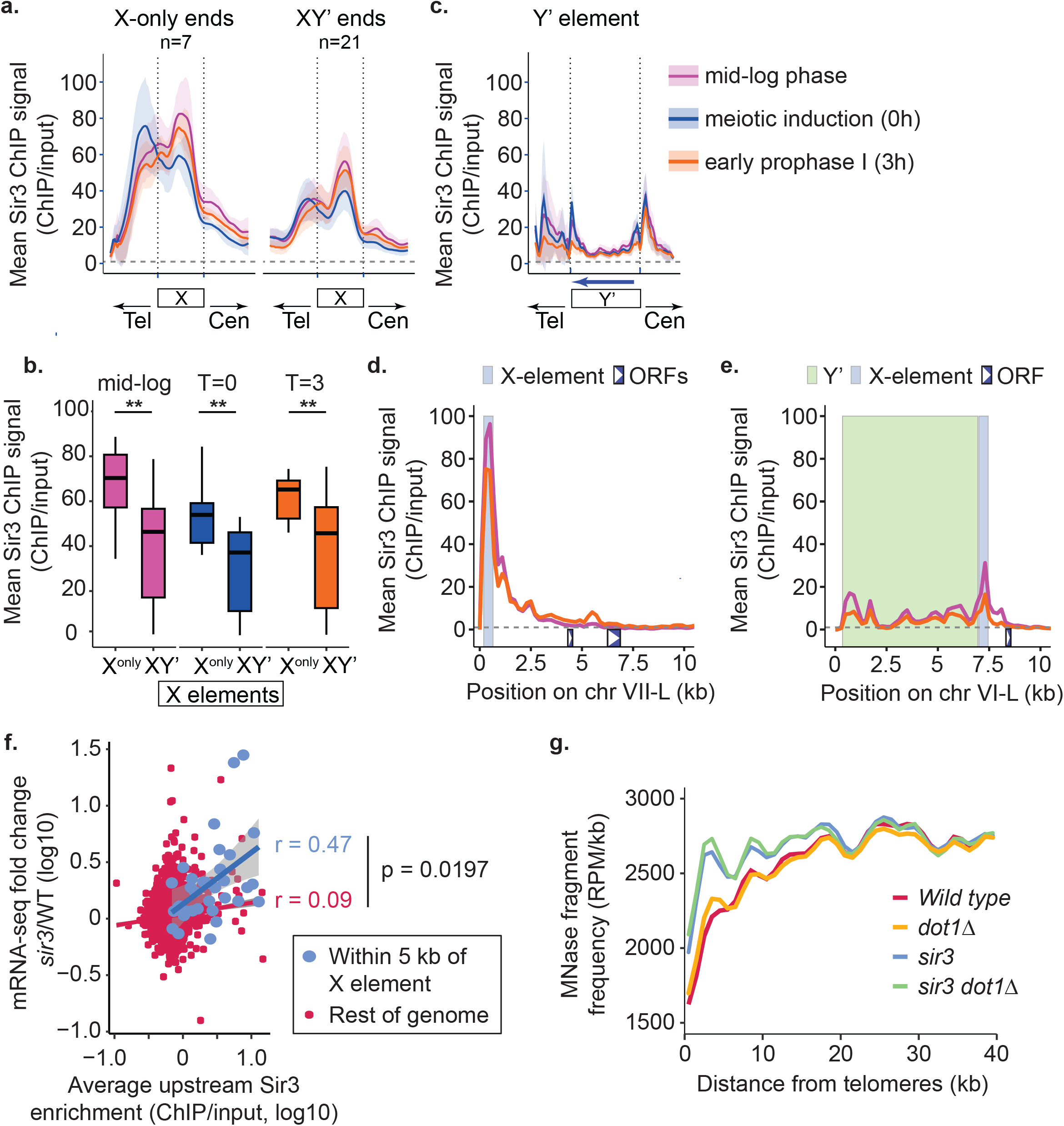
Sir3 occupancy relates to local transcription and MNase accessibility near chromosome ends. **(a)** Metaplots anchored at X elements, stratified by end class, and only fully annotated X elements were used (X-only, n = 7; XY′, n = 21). Flanks scaled to element length (X: 100% each side). Vertical dotted lines mark X boundaries. Shaded bands indicate 95% CIs (see Methods: Metagene analyses, and meta-X and Y’ elements plots). **(b)** Sir3 ChIP signal at X elements on X-only versus XY′ ends during mid-log, meiotic induction (T = 0h), and meiotic prophase (T = 3h). Values represent the mean ChIP/input signal per X element. Box-and-whisker plots show the distribution across elements. Two-sided unpaired Student’s t-tests with BH correction; stars reflect BH-adjusted p (** ≤ 0.01). Statistics (per X element; Cohen’s d; negative = lower at XY′): mid-log (p = 0.007; BH = 0.007; d = −1.22); T = 0h (p = 0.007; BH = 0.007; d = −1.32); T = 3 h (p = 0.002; BH = 0.006; d = −1.11). **(c)** Metaplot anchored at Y′ elements. Fully annotated Y′ only; flanks scaled to 50% of Y′ length. Vertical dotted lines mark Y′ boundaries. Blue arrow indicates Y′-ORF orientation. Shaded bands show 95% CIs (see Methods: Metagene analyses, and meta-X and Y’ elements plots). **(d-e)** Example Sir3 ChIP-seq tracks on chr VII-L and chr VI-L at mid-log and T = 3h. Curves show ChIP/Input. Positions of X elements, Y′ elements and ORFs are indicated (inset white triangle shows ORF orientation). **(f)** mRNA-seq fold change (*sir3*/WT, log10) at T = 3h versus average Sir3 ChIP enrichment in the 250 bp upstream of ORFs. Blue dots represent measurements within 5 kb of X elements, red dots are measurements in the rest of the genome. Pearson r values are shown. P values are based on Fisher’s r-to-z transformation and two-sided z-test. **(g)** MNase-seq fragment frequency (RPM/kb) at T = 3h versus distance from telomeres (0–40 kb) for WT, *dot1Δ*, *sir3*, and *sir3 dot1Δ.* All experiments: averages of two biological replicates.

To determine whether subtelomeric domains with Sir3 spreading exhibit altered transcription of underlying genes, we conducted mRNA-seq analysis on samples collected 3 hours post-meiotic induction in the presence or absence of *SIR3*. Plotting relative fold changes in mRNA levels as a function of the average Sir3 occupancy in the 250 bp upstream of each ORF revealed a significant correlation between Sir3 occupancy and increased mRNA levels upon *SIR3* deletion (**Fig. 7f**). Given the prominent Sir3 peak observed at the start of the Y′ element metaplot (**Fig. 7c**), we also analyzed Y′ element expression 3 hours after meiotic induction. The average Y′ element expression was significantly increased in the absence of *SIR3* (**Supplementary Fig. 11c**). These results indicate that the transcriptional effects of Sir3 are linked to sequences where Sir3 binds, as detected by ChIP-seq.

Sir-dependent transcriptional silencing in vegetative cells is thought to occur by chromatin compaction and promoter occlusion ^52^. MNase-seq analysis of wild type and *sir3* mutants 3h after meiotic induction, revealed a strong Sir3-dependent drop in the relative number of sequenced MNase cleavage fragments in the subtelomeric domains, consistent with chromatin compaction (**Fig. 7g**). Interestingly, the effects on chromatin compaction as measured by MNase-seq extended across more than 20 kb, contrasting with the much more limited extent of detectable Sir3 spreading (**Supplementary Fig. 11a**). These data indicate that Sir3 can influence chromatin compaction even in regions where it is not measurably enriched, possibly through the role of the Sir complex in anchoring telomeres to the nuclear envelope ^53^.

### Sir proteins protect X elements and regions of Sir spreading from meiotic DSBs

We wondered how Sir-dependent heterochromatin (mediated by the Sir2-Sir3-Sir4 complex) and Dot1-dependent axis protein suppression interface with the meiotic DSB machinery near chromosome ends. Genome-wide DSB levels had previously been determined by microarray analysis in *sir2Δ rad50S* strains, but the repetitive nature of the subtelomeric domains had limited coverage, and the X and Y′ elements were not included ^54^. More recently, DSB formation in *sir2Δ* strains was also analyzed using Spo11-oligo sequencing ^19^, which sequences the DNA fragments that remain covalently attached to Spo11 after cleavage ^11^, although telomere-proximal DSB formation was not investigated in that study. Analysis of the Spo11-oligo dataset revealed a significant increase in DSB levels on X elements but not Y′ elements in *sir2Δ* strains (**Fig. 8a**). This pattern mirrors the relative enrichment of Sir3 in these elements, indicating that Sir chromatin suppresses DSB formation in the X elements. In addition, DSB formation was somewhat increased in the subtelomeric domains in *sir2Δ* mutants. Increased DSB induction was correlated with the extent of Sir3 occupancy in wild-type cells (**Supplementary Fig. 12a**). Accordingly, increased DSB formation was also correlated with elevated gene expression in the same regions in *sir3* mutants but not in the rest of the genome (**Supplementary Fig. 12b**). These findings suggest that elevated promoter openness, which enables increased gene expression, also creates a window for meiotic DSB formation.

**Figure 8:**
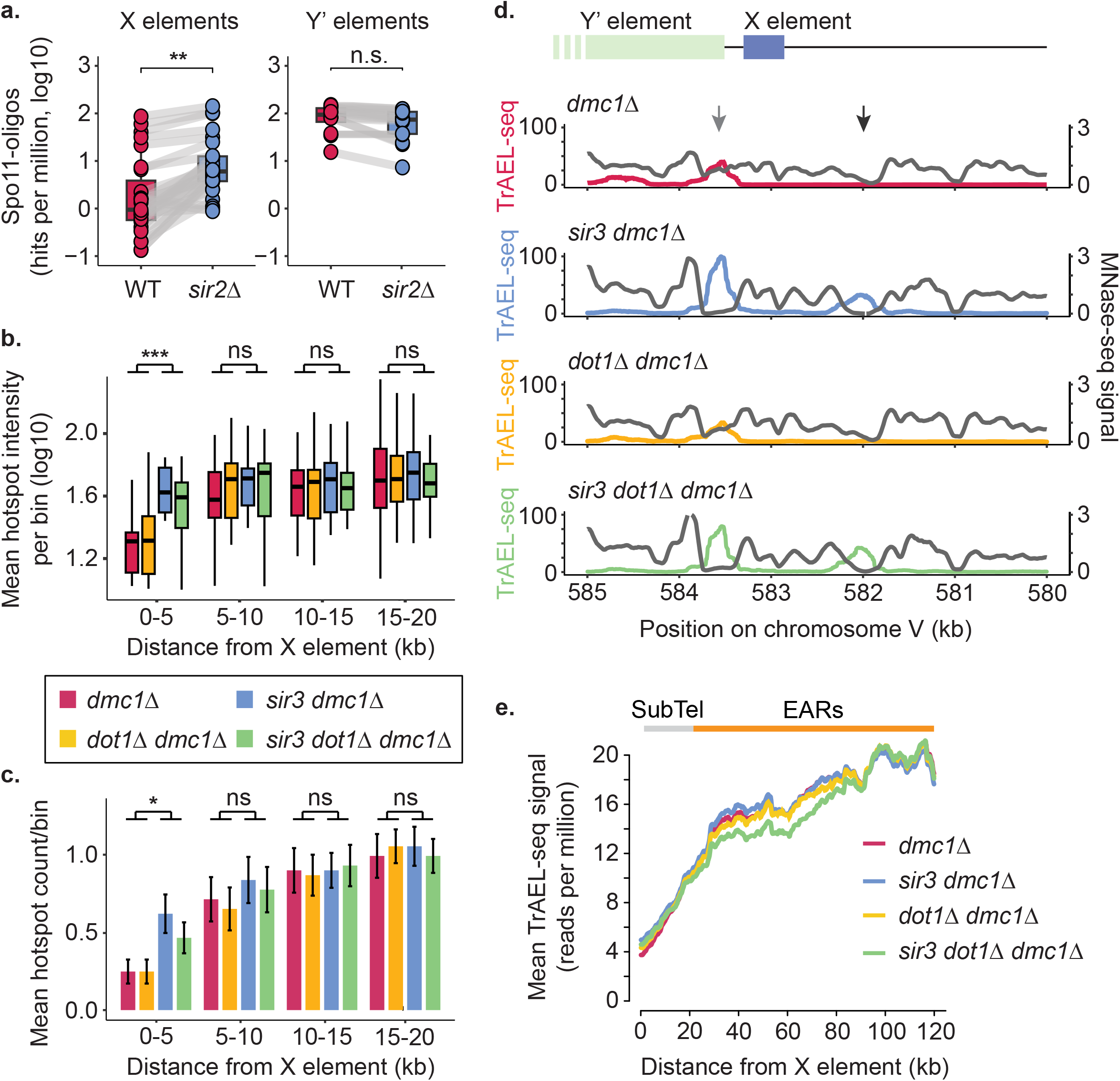
Sir influences DSB formation near chromosome ends. **(a)** Boxplots of log_10_ Spo11-oligo signal (hits per million) at X and Y′ elements in WT and *sir2Δ*, from published datasets ^19,73^. Points are element scores; gray lines connect matched elements across strains. Two-sided Wilcoxon rank-sum tests with BH correction; effect sizes are rank-biserial r (positive = higher in *sir2Δ*). X elements: (p = 7.9x10^-4^, BH = 1.6x10^-3)^; r = 0.83), Y′ elements: (p = 0.127; BH = 0.127; r = 0.32). Stars reflect BH-adjusted p (** ≤ 0.01; n.s., not significant). **(b)** TrAEL-seq hotspot intensity per 5-kb bin as a function of distance from X elements. Log_10_ average peak intensity (RPM) for peaks in each bin in *dmc1Δ*, *sir3 dmc1Δ, dot1Δ dmc1Δ*, and *sir3 dot1Δ dmc1Δ* (see Methods: TrAEL-seq hotspot calling and quantification). Per-bin two-sided Wilcoxon rank-sum tests contrasting (*sir3 dmc1Δ* + *sir3 dot1Δ dmc1Δ*) vs (*dmc1Δ* + *dot1Δ dmc1Δ*), BH-corrected across bins; effect sizes are rank-biserial (positive = higher in *sir3* mutants. Bin 1: p = 3.15 x 10-7; BH = 1.26 x 10-6; r = 0.67. Bin 2: p = 0.237; BH = 0.474; r = 0.13. Bin 3: p = 0.527; BH = 0.703; r = 0.07. Bin 4: p = 0.823; BH = 0.823; r = 0.02. Stars reflect BH-adjusted p (*** ≤ 0.001; n.s., not significant). **(c)** Mean hotspot counts per 5-kb bin as a function of distance from X elements for the indicated strains. Hotspots were identified from TrAEL-seq peak calls. Because most telomere-proximal bins contained only 0-2 hotspots, the data had too few discrete steps for box or violin plots. Therefore, the means of the data are shown as a bar plot (error bars: SEM). Per-bin two-sided Wilcoxon rank-sum tests contrasting (*sir3 dmc1Δ* + sir3 *dot1Δ dmc1Δ*) vs (*dmc1Δ* + *dot1Δ dmc1Δ*), BH-corrected across bins; effect sizes are rank-biserial (positive = higher in *sir3* mutants. Bin 1: raw p = 5.63x10^-3^; BH = 2.25x10^-2^; r = 0.24. Bin 2: raw p = 0.376; BH = 0.752; r = 0.08. Bin 3: raw p = 0.687; BH = 0.915; r = 0.04. Bin 4: raw p = 0.920; BH = 0.920; r = 0.01. Stars reflect BH-adjusted p (* ≤ 0.05; n.s., not significant). **(d)** TrAEL-seq (colored) and MNase-seq (gray) tracks at a representative subtelomeric region (chrV-R) for *dmc1Δ*, *sir3 dmc1Δ, dot1Δ dmc1Δ*, and *sir3 dot1Δ dmc1Δ*. Black arrow indicates a cryptic DSB hotspot that becomes active in the absence of *SIR3*. Gray arrow indicates an unusual Y′-associated hotspot that becomes stronger in the absence of *SIR3.* Other Y′ elements generally do not exhibit altered TrAEL-seq signal. **(e)** TrAEL-seq meta-plots showing mean DSB signal (reads per million) as a function of distance from X elements in the strains indicated. Ranges of subtelomeric domains (gray) and EARs (orange) are indicated above the plot.

To complement and expand this analysis, we analyzed *sir3, dot1Δ*, and *sir3 dot1Δ* mutants by TrAEL-seq, which sequences the exposed 3’ ends that result from Spo11 cleavage ^55^. To avoid signal changes due to DSB repair, this analysis was conducted in a repair-defective *dmc1Δ* background ^56^. We observed several distinct effects on DSB formation near chromosome ends. Consistent with the Spo11–oligo measurements in *sir2Δ* mutants, average hotspot intensity was significantly increased within 5 kb of X elements and the number of significant DSB hotspots nearly doubled in the same regions in strains lacking *SIR3* (**Fig. 8b-c**). Increased DSB formation occurred specifically at sites of increased MNase accessibility (**Fig. 8d**), supporting the model that Sir-dependent DNA occlusion suppresses DSB formation in the heterochromatic parts of the subtelomeric domains. On the other hand, deletion of *DOT1* did not measurably alter DSB formation in the 20 kb from chromosome ends (**Fig. 8b-d**), showing that the increased axis protein deposition in these regions does not translate into higher DSB activity. We also note that overall hotspot activity in the subtelomeric domains remained substantially below genome average in all mutants (**Fig. 8d-e**), indicating that additional layers of regulation contribute to DSB suppression at chromosome ends.

Intriguingly, TrAEL-seq analysis indicated differing genetic interactions between *DOT1* and *SIR3* in other parts of the genome. The most notable large-scale effect was a decrease in DSB levels in the *sir3 dot1Δ* double mutants 30-100 kb from chromosome ends, a genomic range that largely overlaps with the EARs (**Fig 8e**). The reason for this reduction is unclear but may reflect premature downregulation of DSBs in the EARs, which normally experience a longer window of DSB formation ^19^. Further highlighting the different genetic interactions between *DOT1* and *SIR3*, DSB formation around centromeres was decreased to a similar extent in *dot1Δ* and *sir3* single mutants (**Supplementary Fig. 13a**), whereas DSB levels were increased around the ribosomal DNA locus in a Sir3-dependent manner in *dot1Δ* mutants (**Supplementary Fig. 13b**), consistent with previously observed Sir-dependent DSB induction in this region ^57^. As discussed below, the different interactions suggest a combinatorial code that allows these two chromatin regulators to adjust meiotic recombination in a region-specific manner.

## Discussion

Here, we show that key effectors of meiotic recombination, namely axis proteins and the DSB machinery, are suppressed by multiple layers of regulation at chromosome ends. We identified two chromatin regulators, Dot1 and the Sir complex, which contribute to locally suppressing axis protein deposition and DSB formation, respectively. However, additional layers of regulation clearly exist, as DSB levels remain substantially below genome average in *dot1 sir3* double mutants. Additional regulatory layers likely also explain why increases in axis protein binding as observed in *dot1* mutants are not sufficient to increase DSB formation in these regions.

Our data suggest that downregulation of axis protein binding near chromosome ends does not require telomere proximity because placing subtelomeric domains into the chromosome interior of fusion chromosomes did not detectably alter axis protein enrichment. Accordingly, axis protein binding patterns were also remarkably consistent between two different strain backgrounds despite altered distances from the telomeres. These data indicate that the signals that govern the local reduction in axis protein binding are to a large part encoded in *cis.* The molecular nature of these suppressive signals remains to be delineated but our data revealed an intriguing correlation with the reduced coding density of the subtelomeric domains, which tracks remarkably well with the depletion of axis proteins. However, other *cis*-regulatory aspects such as differential replication timing ^33,58^ may also be responsible for the overall lower axis protein levels.

In addition, Dot1 negatively regulates axis protein deposition in X and Y′ elements and the subtelomeric domains. This activity does not require H3K79 and thus either depends on other methylation targets or on Dot1’s methyltransferase-independent activities, several of which have been described ^59,60^. Dot1 ostensibly interferes with Rec8-dependent axis protein recruitment as our analyses suggest that this pathway is specifically weakened near chromosome ends (**Fig. 9**). Given that Rec8 is not significantly depleted near chromosome ends, this interference must specifically impact the interaction between Rec8 and Red1 ^16^.

**Figure 9:**
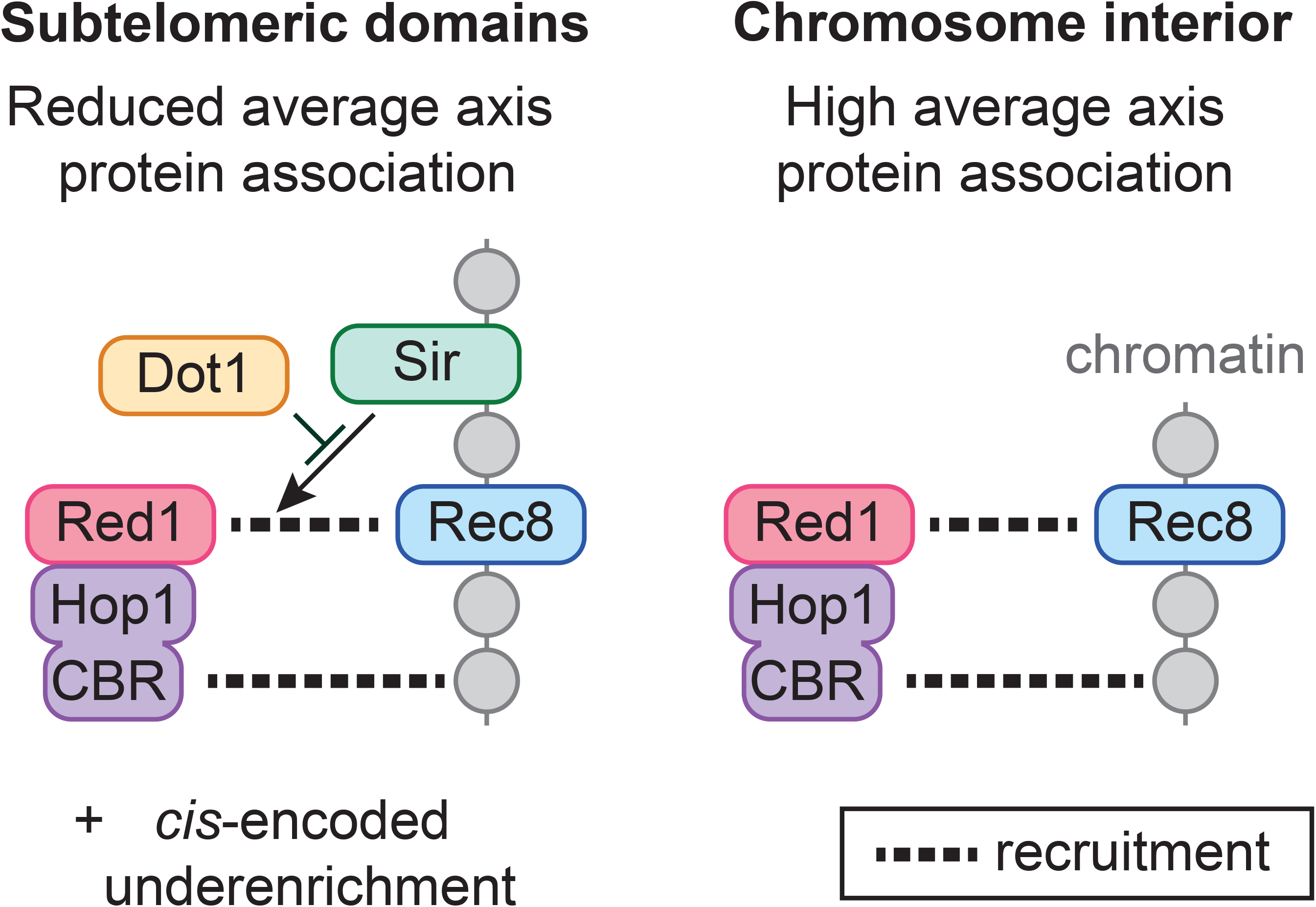
Differential regulation of axis protein deposition near telomeres. Model of how axis protein recruitment is regulated in the chromosome interior (left) and near chromosome ends (right). Axis protein recruitment occurs in parallel through recruitment by Rec8 cohesin (likely via binding to Red1) and through nucleosome binding by the Hop1 chromatin binding region (CBR). Both recruitment pathways are active near telomeres. Dot1 downmodulates Rec8-dependent recruitment, thereby preventing a positive effect of Sir3. In addition, *cis*-encoded features contribute to reduced axis protein binding near telomeres.

Intriguingly, the increased axis protein deposition in *dot1Δ* mutants is rescued by deletion of *SIR3*. This phenotypic epistasis is consistent with Dot1 counteracting the activity of Sir3 to regulate telomere-proximal axis protein deposition. Alternatively, the absence of *DOT1* may create a chromatin environment that allows Sir3 (and by extension the Sir complex) to regulate axis protein binding. This second possibility is supported by the observation that loss of Sir3 (or Sir2 ^19^) alone does not detectably affect the suppression of axis proteins in the subtelomeric domains. An interplay between the Sir complex and Dot1 was previously shown to affect checkpoint regulation during meiotic recombination, although we note that in that situation, loss of *SIR2* and *DOT1* both inactivate checkpoint activity possibly because of a redistribution of Sir2 away from sites of checkpoint activity in *dot1* mutants ^61^. In vegetative cells, Dot1 helps constrain Sir-protein dependent silencing both through H3K79 trimethylation and by competing for the same nucleosomal binding site ^62^. This two-pronged mechanism may explain why mutation of H3K79 was not sufficient to recapitulate the increased axis protein deposition seen in *dot1Δ* mutants. Interestingly, even though the axis proteins act as recruitment platforms for the DSB machinery ^12^, increased axis protein deposition was not sufficient to increase DSB formation in the telomere-proximal sequences in *dot1Δ* mutants, indicating that additional layers of regulation suppress DSB formation in these regions.

In addition to its role in regulating axis proteins, we show that the Sir complex suppresses DSB formation at a small number of subtelomeric hotspots through local chromatin compaction of gene promoters, the preferred DNA substrates for DSB formation in yeast ^11^. Our data strongly imply that DSB suppression is a direct consequence of the enrichment of Sir3 in these regions. Furthermore, the strict spatial overlap between increased MNase sensitivity and sites of DSB formation in *sir3* mutants supports the notion that nucleosomes interfere with meiotic DSB formation, presumably by restricting access to the DNA substrate ^11,45,63^. However, our data also show that the increased MNase sensitivity in *sir3* mutants extends across more than 20 kb from telomeres, whereas hotspot activation is much more spatially restricted. These data indicate that increased DNA accessibility alone is in most cases not sufficient to activate meiotic DSB formation in the subtelomeric domains.

While this study focused on the subtelomeric domains, our genome-wide data also showed effects of Dot1 and the Sir complex in other parts of the genome, including the EARs, the pericentromeres, and the rDNA-flanking regions. H3K79me3 is abundantly present in those regions, and Dot1’s distribution in vegetative cells closely matches H3K79me3 ^64^, suggesting that Dot1 and H3K79me3 may directly influence axis protein deposition and/or DSB formation in some of those regions. By contrast, Sir3 is not noticeably enriched in the EARs or the pericentromeres in our analyses, although genome-wide effects of *sir* mutation on meiotic DSB formation were also noted previously ^54^. It is possible that the Sir complex levels in these regions are too low to be measured confidently by our methods. Alternatively, some effects may be indirect as both the Sir complex and Dot1 are transcriptional regulators and thus could change the expression levels of key meiotic regulators to modulate regional axis protein deposition or DSB activity. Such indirect effects may also contribute to the varying genetic interactions between the Sir complex and Dot1 observed in different genomic regions.

Chromosome ends differ in their sequence composition, spreading of the Sir complex, and the location of genes, DSB hotspots, and axis binding sites, creating inherently chromosome end-specific variability in the range and presence of individual suppressive effects. Nevertheless, downregulation of axis protein deposition and DSB induction within about 20 kb from chromosome ends is remarkably uniform (**Supplementary Fig. 3a-b**). Thus, an overall selective pressure likely exists to reduce axis protein binding and DSB induction at chromosome ends, and multiple molecular mechanisms have the potential to satisfy this goal.

One selective pressure near chromosome ends is the need to suppress meiotic crossovers. Telomere-proximal crossovers are less effective at maintaining linkages between homologous chromosomes ^65^ and are associated with increased risk of Down syndrome in humans ^23,24^. Axis proteins, in particular, are a logical target for such suppression, as they not only recruit DSB factors but also help target DSB repair to homologous chromosomes by preventing sister chromatid recombination ^66^. Favoring repair from sister chromatids may also reduce the risk of non-allelic recombination events. Chromosome ends are especially prone to non-allelic homologous recombination because of their repetitive sequences and the close physical proximity of telomeres along the nuclear envelope during meiotic prophase ^21,67,68^. Having multiple regulatory layers independently suppress axis protein deposition and DSB formation may thus establish a robust protective mechanism to shield chromosome ends from non-allelic and unproductive recombination and ensure proper meiotic chromosome segregation.

## Methods and Materials

### Yeast strains and growth conditions

All the strains utilized in this study belonged to the SK1 background, except for the analysis using the published fusion chromosome strains, which were SK1/S288c hybrids, and the spike-in analysis, which used meiosis-competent S288c (SK288c) strains (see Methods: Spike-in Normalization). The genotypes are detailed in **Supplementary Table 1.** For experiments using vegetative cells, cultures were grown in YPD medium overnight at room temperature until saturation. The following day, the cells were diluted to an OD_600_ of 0.2 and incubated at 30°C until they reached an OD_600_ of 1. At this stage, 50 ml of culture was collected for ChIP-seq.

To induce synchronous meiotic cultures, strains were grown for 24 hours in YPD medium at room temperature. Cells were enriched in the G1 phase by inoculating them at OD_600_ = 0.3 in pre-sporulation BYTA medium (1% yeast extract, 2% bactotryptone, 1% potassium acetate, 50 mM potassium phthalate) for 16.5 hours at 30°C. Cells were washed twice with sterile water and transferred into SPO medium at OD_600_ = 1.9 (SPO: 0.3% potassium acetate, 0.001% acetic acid). Cultures were incubated at 30°C on a shaker, and the time of inoculation into SPO was defined as T = 0 hours. At the specified timepoints, 25 ml of culture was collected for ChIP-seq and 1.6 ml for mRNA-seq. The synchrony of meiotic cultures was validated using flow cytometry of DNA content.

### Chromatin immunoprecipitation (ChIP) & sequencing

Cells collected at the indicated timepoints (T = 3h unless noted otherwise) were pelleted and immediately crosslinked for 30 minutes in a 1% formaldehyde solution at room temperature with gentle shaking. Subsequently, the crosslinking reaction was quenched by incubating the cells for 5 minutes at room temperature with 125 mM glycine. Following this, the cells were processed according to the protocol outlined in ^16^ and immunoprecipitated with 2.5 μl of either anti-Sir3 (HM01065, kind gift of S. P. Bell), anti-Red1 (#16440, kind gift of N. Hollingsworth) or anti-H3K79me3 (abcam, ab2621). For spike-in normalization (SNP-ChIP ^36^), SK288c cross-linked meiotic samples were added to respective samples at 20% prior to ChIP processing. Library preparation, quality, and quantity checks were then performed as described in ^15^. All the prepared chromatin immunoprecipitation (ChIP & SNP-ChIP) libraries were sequenced as 150-bp paired-end reads on Illumina NextSeq 500 or Element Biosciences’ Aviti instruments. The sequencing runs were conducted by the Genomics Core at New York University Center for Genomics and Systems Biology.

### Processing ChIP-seq reads from sequencing

Sequencing reads were aligned to the SK1 genome ^35^ using Bowtie 2 (Version 2.4.2) ^69^. To improve mapping of reads to repetitive subtelomeric domains, the Bowtie 2 read reporting mode was configured to allow multiple alignments, reporting the best match. MACS2 2.1.1 was used to extend reads in the 5’-> 3’ direction to a final length of 200 nt. SPMR (signal per million reads) normalization was applied to both input and ChIP pileups using MACS2. The resulting fold-enrichment files were used for downstream analysis in R. The ChIP-seq pipeline and analysis scripts are available on the Hochwagen Lab GitHub page https://github.com/hochwagenlab/ChromosomeEnds.git.

### mRNA-sequencing & analysis

mRNA was extracted from cells collected at T = 3h. mRNA extraction, first-and second-strand synthesis, and library preparation were conducted following the procedures outlined in ^70^. The resulting libraries were sequenced as 150-bp paired-end reads on an Illumina NextSeq 500 instrument. The sequencing run was executed by the Genomics Core at New York University Center for Genomics and Systems Biology. The GTF (General Transfer Format) file was modified to include all completely annotated Y′ elements. Subsequently, reads obtained from Illumina sequencing were aligned to the SK1 genome ^35^, and the counts mapping to the modified SK1 gtf file were determined using the nf-core RNA-Seq pipeline ^71^. The relative abundances, measured as transcripts per million (TPM) in Salmon ^72^, were used for downstream analysis in R.

### Spo11-oligo mapping

Published Spo11-oligo datasets from WT and *sir2Δ* strains ^19,73^ were analyzed. Adaptors were removed from the reads using fastx_clipper from the FASTX Toolkit (version 0.0.14) and reads shorter than 15 nt were discarded. The clipped reads were aligned to the SK1 genome ^35^ using Bowtie 2 (Version 2.4.2) ^69^. The read alignment mode was set to local, and the reporting mode allowed multiple read alignments while reporting the best alignment. SPMR (signal per million reads) normalization was performed on the read pileups.

### Mononucleosomal DNA preparation

For MNase-seq, 50 ml of synchronized meiotic cultures was harvested at T = 3h and crosslinked with 1% formaldehyde for 30 minutes at room temperature. Crosslinking was quenched with 125 mM glycine, and cells were spheroplasted in Buffer Z (0.5 M sorbitol, 50 mM Tris-HCl pH 7.4, 10 mM β-mercaptoethanol) containing zymolyase T100. Spheroplasts were pelleted and resuspended in NP buffer (1 M sorbitol, 50 mM NaCl, 10 mM Tris-HCl pH 7.4, 5 mM MgCl₂, 1 mM CaCl₂, 0.5 mM spermidine, 1 mM β-mercaptoethanol, 0.075% NP-40) and digested with 5–80 units of micrococcal nuclease (MNase; Thermo Fisher Scientific, EN0181). MNase was diluted from a 300 U/µl stock to 20 U/µl, and mononucleosome-sized fragments were consistently obtained using 2 µl of a 1:4 dilution of the 20 U/ µl stock (final concentration 5U/µl; 10 units in total). Digestions were stopped with SDS and EDTA, followed by proteinase K treatment and overnight incubation at 65°C. Subsequent steps, including DNA purification and library preparation, were performed as described in ^11^. Libraries were sequenced as 150-bp paired-end reads on an Illumina NextSeq 500, and the sequencing run was carried out by the Genomics Core at the New York University Center for Genomics and Systems Biology.

### MNase-seq data analysis

To improve read alignment in repetitive subtelomeric domains, Bowtie 2 was configured to report multiple alignments using the “best match” mode as described for ChIP-seq. For signal quantification, fragments were processed with MACS2 (--nomodel --keep-dup all --SPMR) to generate normalized coverage tracks. To map MNase cleavage sites at nucleotide resolution, the 5′ end of each properly paired fragment was extracted from BAM files. This yielded single-base BED files representing MNase cut frequency, which were used for downstream metaplot analysis of chromatin accessibility.

### Plug preparation for TrAEL-seq

Plugs were prepared from meiotic cultures harvested at T = 5h to allow for accumulation of DSBs in the *dmc1* mutant background. For each strain, 20 ml of culture was pelleted and washed twice with CHEF TE buffer (10 mM Tris-HCl, pH 7.5, 50 mM EDTA, pH 8.0). The cell pellet was resuspended in 300 µl CHEF TE, followed by the addition of 4 µl zymolyase T100 (10 mg/ml). The mixture was briefly vortexed and incubated at 42°C for approximately 30 seconds. Low-melting-point agarose (1% SeaPlaque GTG in 125 mM EDTA, pH 8.0), prewarmed to 42°C, was added and transferred into plug molds. Plugs were solidified on ice for 10 minutes before being transferred into LET buffer (10 mM Tris-HCl, pH 7.5; 0.5 M EDTA, pH 8.0). Plugs were incubated overnight at 37°C. The following day, plugs were treated with proteinase K in NDS buffer (10 mM Tris-HCl, pH 7.5; 0.5 M EDTA; 1% N-lauroylsarcosine) at 50°C overnight. After digestion, plugs were washed sequentially: first with CHEF TE for 1 hour at room temperature, followed by two washes with freshly prepared ice-cold CHEF TE containing 1 mM PMSF (from 100 mM stock in isopropanol), with each wash incubated for 1 hour in the cold room. A fourth wash was performed with RNase T1 in CHEF TE, incubated at 37°C for 1 hour, followed by a final wash with CHEF TE at room temperature for 1 hour. Plugs were stored at 4°C in CHEF TE until further use.

### TrAEL-seq Library Preparation, Sequencing, and Data Analysis

TrAEL-seq library preparation, sequencing and data analysis were conducted following the procedures outlined in ^55^. TrAEL-seq signal pileups were generated using MACS2 (v2.2) and normalized pileups were generated using SPMR normalization. Downstream analyses were done in R. We note that telomeres can have free 3’ ends that lead to a Spo11-independent TrAEL-seq signal. Because this signal showed strong batch effects that impacted the nearby Y′ and X elements, we were unable to confidently analyze DSB formation in Y′ and X elements using this method.

### Statistical data analysis

#### 1) Spike-in normalization

To allow for quantitative comparisons of chromatin immunoprecipitation (ChIP) enrichment between different strains or conditions, we performed spike-in normalization using SNP-ChIP ^36^. A known amount of SK288c chromatin (S288c strains engineered to efficiently enter meiosis) was used as a spike-in control, with sequence reads mapping to the S288c genome serving as a reference for normalizing sequencing depth and ChIP efficiency. The ratio of SK1 reads to S288c reads was then used to scale the enrichment values, allowing for direct comparison of protein abundance across different samples. This method is particularly useful for quantifying relative changes in protein occupancy between mutant and wild-type strains. Data sets from single strains, such as the analysis of wild-type strains presented in **Fig. 1**, were analyzed without spike-in normalization, as no common denominator was necessary for these analyses.

#### 2) Distance from telomeres plots

The x-axis represents the distance from the annotated telomeric repeats, beginning at telomeres and extending inward. The y-axis indicates signal intensity, normalized so that the genome-wide average equals 1, or spike-in normalized values where applicable. Data were processed as in ^19^ by binning signals into consecutive 200-bp windows, followed by kernel smoothing to minimize noise and highlight underlying trends.

#### 3) Bootstrapping plots

Violin plots represent bootstrapped distributions obtained by performing 1000 iterations of resampling, where, for each iteration, 32 regions of 20 kb (or 16 regions for centromeric analyses) were sampled with replacement. The violin plots display the resulting distributions, indicating the median values along with two-sided 95% confidence intervals. Two-sided empirical p-values were computed against this distribution and BH-corrected. Effect sizes were measured via Cohen’s d,

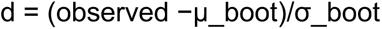

where observed is the mean fold-enrichment in the focal regions; μ_boot is the mean of the bootstrap distribution of means; and σ_boot is the standard deviation of that bootstrap distribution.

#### 4) Metagene analyses, and meta-X and Y’ elements plots

Only fully annotated X/Y′ elements were analyzed; partially annotated copies were excluded because boundary-anchored profiles required accurate start and end coordinates. The excluded partial X elements are indicated with stars in **Supplementary Fig. 2** (coordinates: VIII:9643-9820, X:15487-15645, XII:1020084-1020314, XV:1047720-1047878, XVI:9407-9565). Metagene analyses followed previously published approaches ^13^. Similarly, to align variable-length X and Y’ elements, we proportionally extended flanking regions based on the element length (100% extension for X elements and 50% for Y’ elements, as indicated in the figure legends). Each extended region was then divided into 100 equal bins, scaling each element to fixed relative positions: X elements occupied the middle third (bins 33–66), and Y’ elements occupied the middle half (bins 25–75). Signal intensities were averaged across these bins, ensuring proportional alignment at element boundaries (e.g., bin 33 corresponded to the start of X elements, bin 25 corresponded to the start of Y’ elements).

#### 5) Confidence bands vs. bootstrapping

We used different approaches to determine significance depending on the structure of the underlying data. For metaplots of gene bodies, X, or Y′ elements, the structure of the underlying sequences, and hence the distribution of chromatin marks and axis proteins, was relatively consistent after adjusting for the length of the elements. This structure allowed meaningful confidence bands to be shown. By contrast, the distribution of axis binding sites was inherently sparse and differed between chromosome ends. As a result, a confidence band was not meaningful as the differences in the positions of axis sites would drown out consistent differences in signal across ends. To circumvent this problem, we instead combined the signal of the entire region and evaluated how significantly different the mean was from the rest of the chromosome using bootstrapping.

#### 6) Quantification of Red1 and Rec8 signals

Rec8 peaks in **Fig. 4a** were derived from Rec8 ChIP-seq data in wild-type cells (the same dataset used in **Fig. 1b-f**). For each Rec8 peak region, the Red1 signal was calculated as the average score across the entire Rec8 peak interval. Peaks were annotated as centromere-proximal (within ±10 kb of a centromere), telomere-proximal (within 20 kb of a chromosome end), or non-centromere/non-telomere to contextualize their genomic distribution. If peak regions were straddling the boundaries, they were included as centromere-proximal/telomere-proximal. Straddling Peaks: centromeres 0.16% and telomeres 0.08% of total peaks. We used Rec8 peaks because the Rec8 signal did not change significantly compared to genome average in the subtelomeric domains. The reduction of Red1 signal near telomeres meant that some sites that have abundant Rec8 binding near telomeres did not have sufficient Red1 binding to be called as peaks.

#### 7) Placing peaks at chromosome ends into register

Peaks were called independently in each genome as local signal maxima at least 1.3-fold above the genome-average signal, spaced at least 3 kb apart. Profiles were stacked using X elements (where annotated) or telomeres and tested at a range of relative shifts, both toward the telomere and toward the chromosome interior. Whichever position brought the most SK1 and S288c peaks into close correspondence was chosen as the final alignment. Alignment quality was confirmed by whole-profile and peak-region correlation, together with visual inspection of each aligned profile.

#### 8) Quantification of Sir3 spreading from chromosome ends

Sir3 spreading was quantified using a custom R pipeline. For each condition, ChIP-seq fold-enrichment over input (FE) signals were extracted from chromosome-end regions from the X element toward the centromere. These regions were divided into consecutive 250 bp bins, and the average FE score was calculated for each bin. A dynamic threshold was defined as the upper bound of the 95% confidence interval of genome-wide FE values. Spreading was identified starting at the X-element boundary as the first continuous stretch of bins with FE scores above this threshold. The spreading region was considered to end when two consecutive bins fell below the threshold; the final bin above threshold before this drop marked the endpoint of Sir3 spreading. Spreading distances were aggregated across chromosome ends and visualized using violin plots.

#### 9) Promoter correlation plots

Promoters were defined as the 250-bp window immediately upstream of annotated genes. Spo11-oligo signal was HPM-normalized and averaged per promoter in WT and *sir2Δ*. We plotted log_10_(*sir2Δ*/WT). To avoid inflated ratios from near-zero denominators, we set a data-driven floor at the 2^nd^ percentile of promoter means pooled across WT and *sir2Δ* and excluded promoters only when both WT and *sir2Δ* means were below this floor (117 out of 5541; 2.11%). Results were robust to using a 5% floor (correlations changed only minimally). After filtering, the plotted set contained 34 genes within 5 kb of an X element (blue) and 5,356 other genes (red). The same promoter set was used for **Fig. 7f** and **Supplementary Fig. 12a–b**. RNA-seq fold changes were obtained with DESeq2 (two biological replicates per condition) and are shown as log_10_. Sir3 ChIP was genome-normalized (ChIP/Input). Values are means of two independent biological replicates and were reproducible between replicates.

#### 10) Hotspot calling in TrAEL-seq data

DSB hotspots were identified from TrAEL-seq data using MACS2 (narrowPeak mode) on pooled biological replicates for each genotype, with the q-value (FDR) threshold set to control the false discovery rate. Hotspot totals were: *dmc1Δ* (n = 3,242), *sir3 dmc1Δ* (n = 3,224), *dot1Δ dmc1Δ* (n = 3,182), and *sir3 dot1Δ dmc1Δ* (n = 3,130). For quantification, signal was taken from MACS2 treat_pileup bedGraphs (RPM) and summarized as the mean RPM across each hotspot interval. For each telomere, the 0–20 kb region inward of the annotated X-element was tiled into 5-kb bins (bins 1–4 from the X element). Each hotspot was assigned to exactly one bin using its start coordinate, preventing double-counting of hotspots that span multiple bins.

## Supporting information

Supplementary Figures and Tables are attached in a word document.

## Data reporting

The datasets generated and analyzed in this paper, excluding published datasets, have been deposited in the Gene Expression Omnibus (GEO). The datasets can be accessed with the accession numbers GSE287129 (ChIP-seq), GSE287127 (mRNA-seq), and GSE287130 (TrAEL-seq). Additionally, all ChIP-seq datasets used in this study are listed in **Supplementary Table 2**, and all Spo11-oligo, mRNA-seq, and TrAEL-seq datasets are listed in **Supplementary Table 3**.

## Acknowledgements

We thank Stephen P. Bell for generously providing the Sir3 antibody, Nancy Hollingsworth for the Red1 antibody, and Akira Shinohara for the *hht1/2-K79R* strains. We also acknowledge the Genomics Core at the New York University Center for Genomics and Systems Biology for their valuable technical assistance and expertise in data processing. This work was supported in part by the NYU IT High-Performance Computing resources, services, and staff expertise. We are grateful to the Zegar Family Foundation for their generous support. TrAEL-seq library sequencing and processing were performed by the Genomics (Geno06) and Bioinformatics (Bioinf01) teams at the Babraham Institute, which receive financial support from the Institute Core Capability Grant (BBSRC CCG). This research was financially supported as part of NIH grant R35 GM148223 to A.H. A.R.R. acknowledges support from a Fleur Strand Graduate Fellowship from the Department of Biology, as well as a Henry MacCracken Fellowship and a Dean’s Dissertation Fellowship from the NYU Graduate School of Arts and Science. J.H. acknowledges funding from the BBSRC (BI Epigenetics ISP; BBS/E/B/000C0523), and K.M. acknowledges funding from the BBSRC (BB/W509917/1). The funders had no role in the preparation of this manuscript.

## Author contributions

Conceptualization - A.R.R., V.V.S., H.G.B., and A.H; Investigation & Formal analysis - A.R.R., K.M., V.V.S., H.G.B., N.J.P., J.H., A.H; Computational Pipeline development - A.R.R.; Manuscript writing (initial draft) - A.R.R. and A.H.; Manuscript editing - all authors.

## Supplementary Figures

**Supplementary Figure 1.**
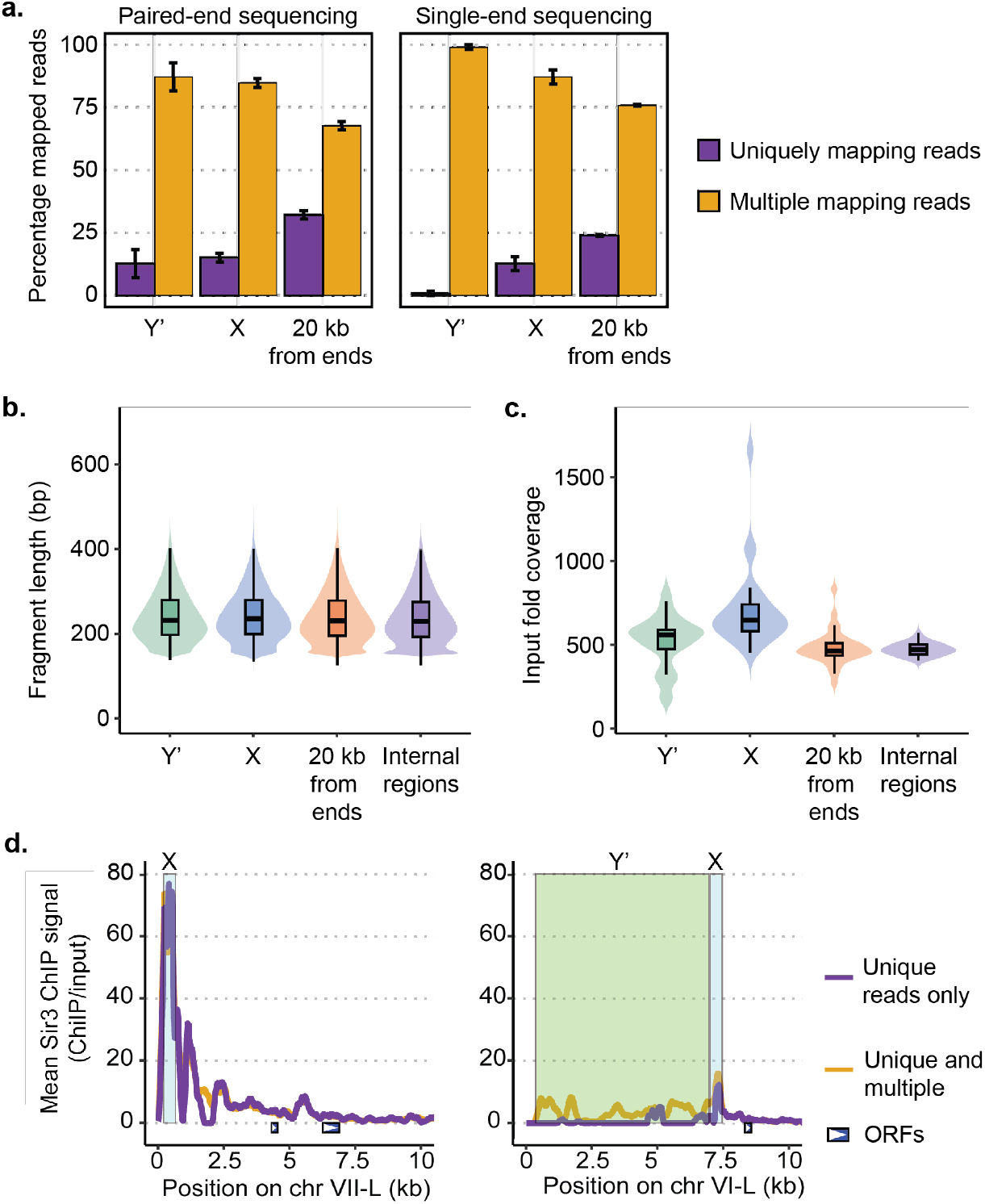
: Enhanced coverage at chromosome ends with a tailored mapping workflow. **(a)** Fraction of uniquely mapping and multiple-mapping reads within Y′ elements, X elements, and the terminal 20-kb (subtelomeric domains), shown for paired-end and single-end libraries. Bars show mean across replicates; error bars, SD. **(b)** Fragment-length distributions for unbiased input DNA in Y′, X, terminal 20 kb, and internal regions. Violin plots show the full distribution of paired fragments. Embedded boxplots mark the median and interquartile range **(c)** Input fold coverage across the same region classes, displayed as violin plots with embedded boxplots. For X, Y′, and terminal 20-kb regions, coverage was calculated as the averaged depth per element. For internal regions, we averaged depth within non-overlapping 20-kb windows defined after excluding X, Y′, and terminal 20-kb regions. The increased input signal for X elements likely reflects the missing X-element annotations on several chromosomes (**Supplementary** Fig. 2). Since all ChIP profiles are plotted as ChIP/input, any increase in input signal will be mirrored in the ChIP signal and thus will not affect conclusions. **(d)** Sir3 ChIP profiles at representative subtelomeres (chr VII-L; chr VI-L), comparing unique versus unique+multi-mapping pipelines. Shaded rectangles mark X and Y′ elements. Blue boxes and arrowheads indicate ORFs and their directions. Values are means of two independent biological replicates and were reproducible across replicates.

**Supplementary Figure 2.**
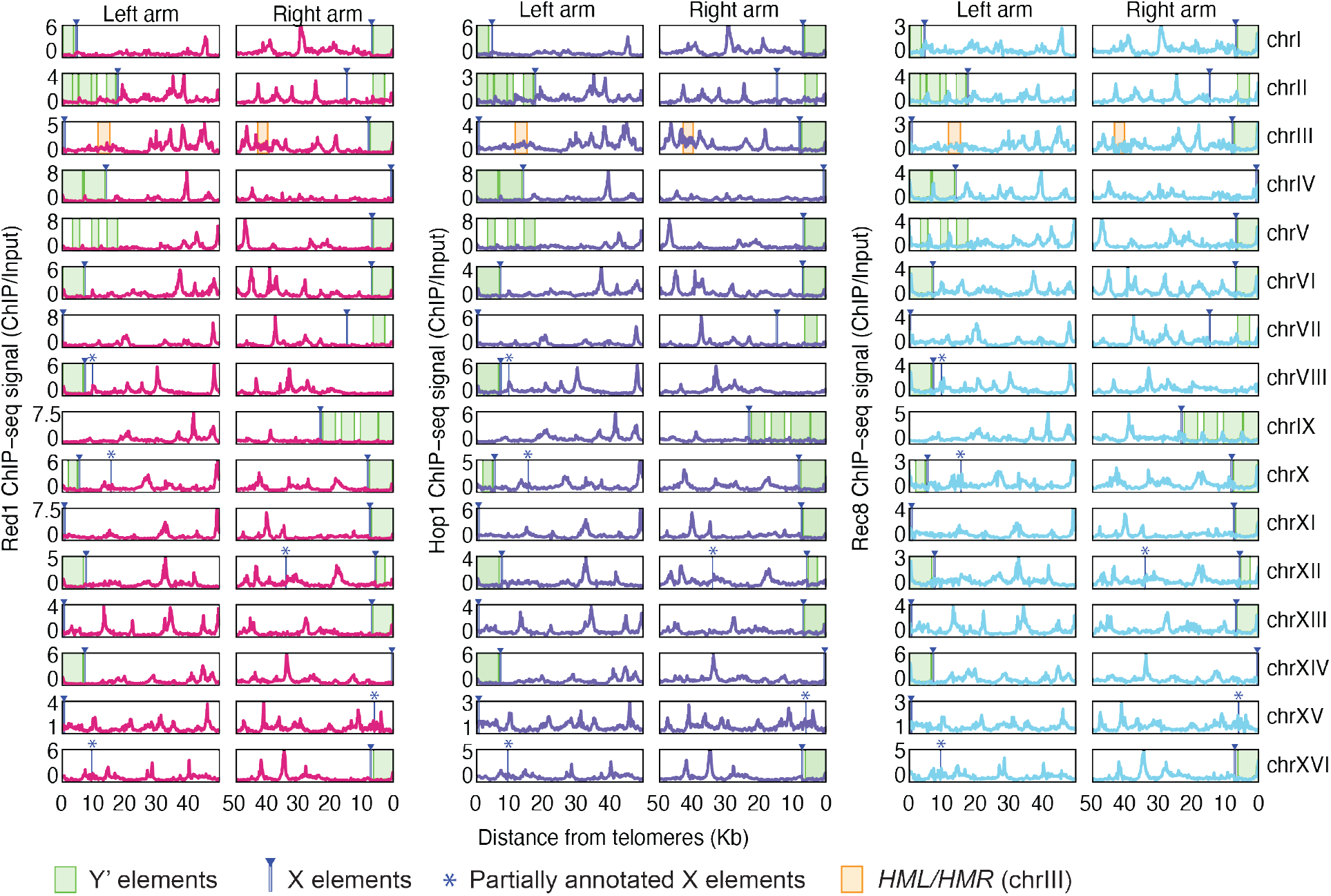
: Axis-protein ChIP profiles across all chromosome ends. Distance-from-telomere ChIP/Input signal for Red1, Hop1, and Rec8 at every left and right arm (32 ends) in WT early prophase I (T = 3h). Green blocks mark all annotated Y′ elements and blue blocks with arrowheads mark all annotated X-elements (fully and partially annotated copies shown: X, n = 32; Y′, n = 31). The partially annotated copies (marked with stars) are located more internally and may reflect degenerating X elements or incorrect annotation due to incomplete sequence conservation and the small size of X elements. These partially annotated X elements were not used for analyses shown in this study. Signals are from published datasets ^1-3^. The positions of the silent mating type loci near the left (*HML*) and right (*HMR*) telomeres of chr III are indicated as orange boxes.

**Supplementary Figure 3.**
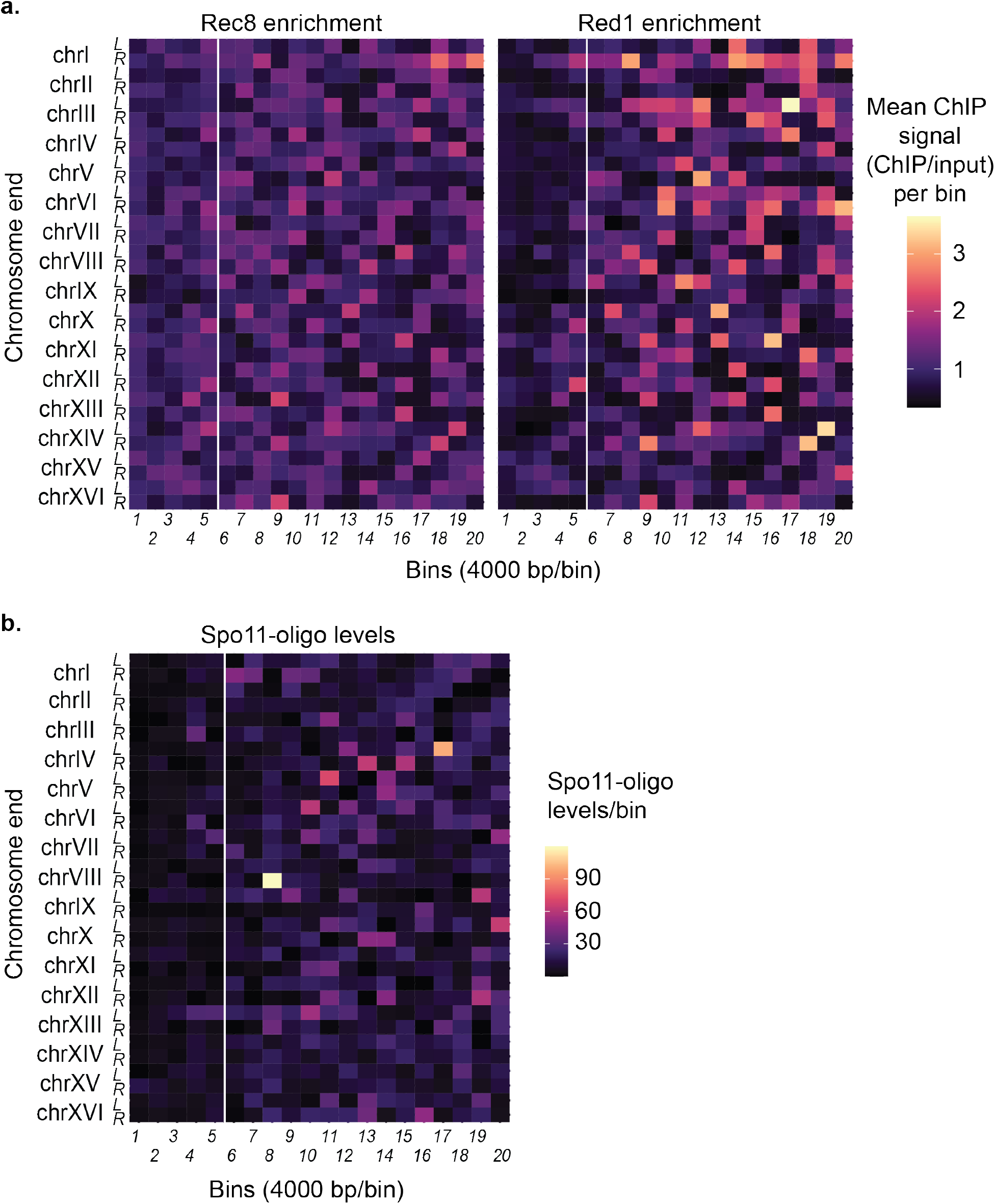
: Consistent axis-protein and Spo11-oligo depletion across chromosome ends. **(a)** Heatmaps of Rec8 and Red1 ChIP enrichment (ChIP/Input) in WT early prophase I (T = 3h) from published data ^2,3^. Each row is a chromosome end (left/right indicated). Columns are contiguous 4-kb bins extending inwards from the telomeres. Colors show the mean signal per bin; the vertical white line marks 20 kb from the end. **(b)** Heatmap of Spo11-oligo density (WT; data from ^4^) in the same 4-kb binning method. Values are means of two independent biological replicates and were reproducible across replicates.

**Supplementary Figure 4.**
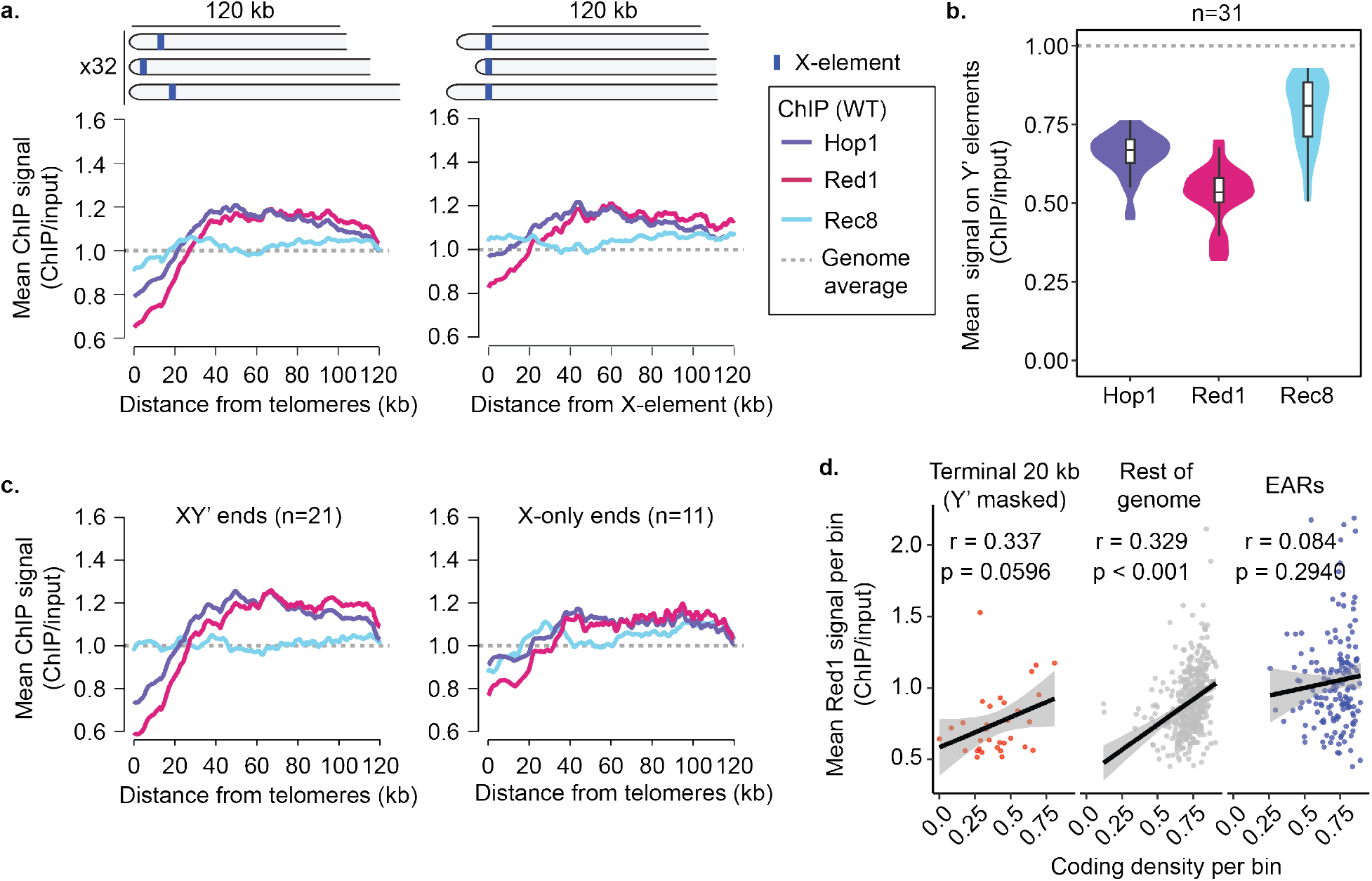
: Low axis-protein enrichment on Y′ elements contributes to the depletion seen in metaplots. **(a)** Mean enrichment of Hop1, Red1, and Rec8 (ChIP/Input) in WT early prophase I (T = 3h) from published datasets ^1-3^, normalized to a genome average of 1 (gray dashed line). Left panel: distance from telomeres; right panel: distance from the annotated X-element anchor. Curves were computed as described in Methods: Distance-from-telomere profiling. **(b)** Mean enrichment of Hop1, Red1, and Rec8 on Y′ elements (n = 31), normalized to a genome average of 1 (gray dashed line). Violin plots show the full distribution of means. Embedded boxplots mark the median and interquartile range. **(c)** Distance-from-telomere profiles stratified by end class: XY′ ends (n = 21) versus X-only ends (n = 11) for Hop1, Red1, and Rec8. **(d)** Relationship between coding density and mean Red1 enrichment. Scatter plots show non-overlapping 20-kb bins at dots. Black lines are least-squares fits, with 95% CIs shown in gray. Pearson r values are indicated. P values are based on two-sided t-tests on the slope. “Y′ masked” means Y′ sequence is excluded from both measurements (we mask Y′ bases when tallying coding density and when averaging Red1; bins that are entirely Y′ are excluded). Including the Y′ elements eliminated the correlation in the last 20 kb (r = -0.174; p = 0.341).

**Supplementary Figure 5.**
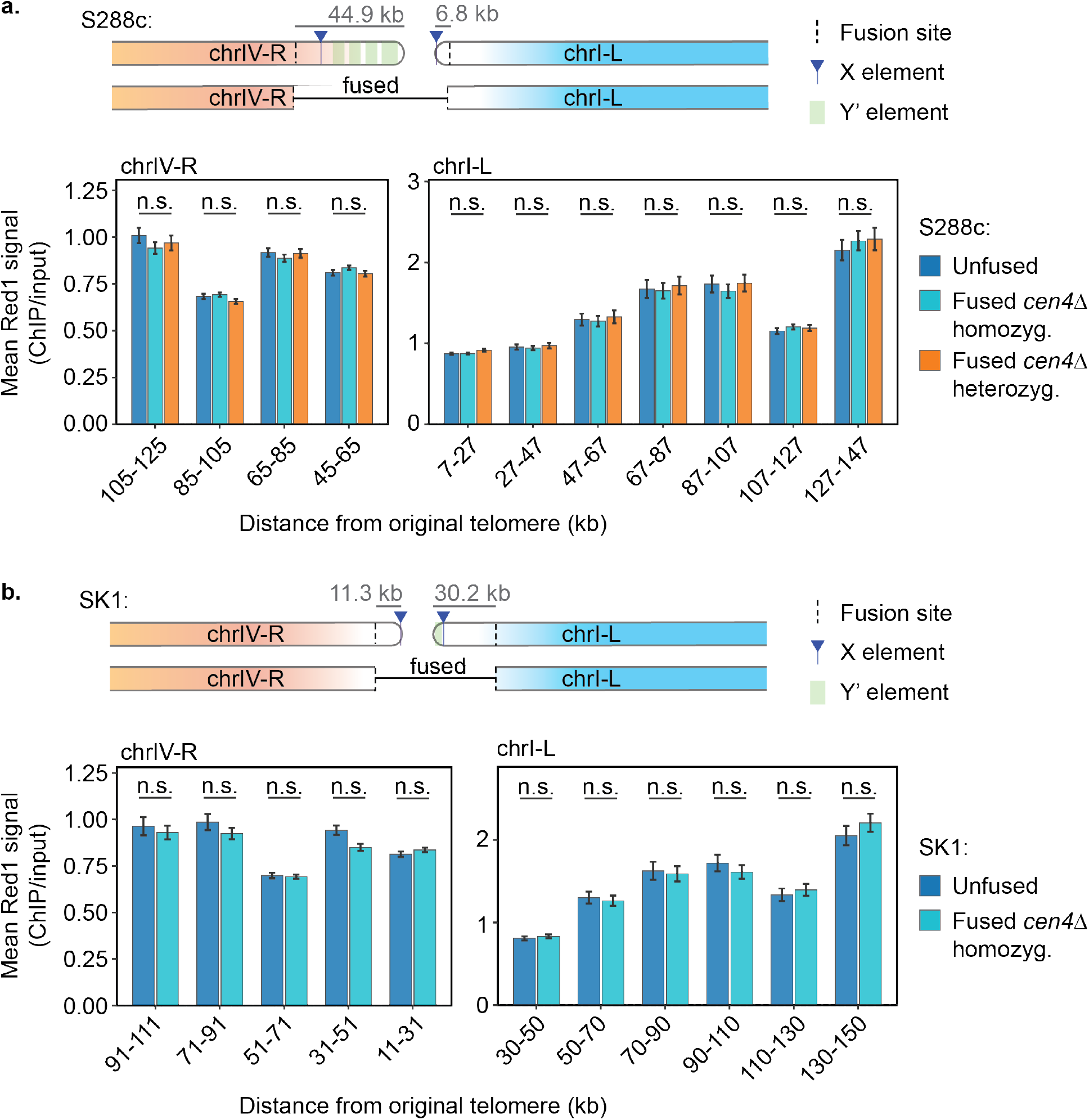
: No effect of chromosome fusion on Red1 enrichment near site of fusion. Mean Red1 enrichment (ChIP/Input) in 20-kb bins tiling from the fusion sites on wild-type and fused chromosomes IV and I from S288c (**a**) or SK1 (**b**). Schematics indicate which regions were deleted axs part of the fusion process. Because of the effects of centromeres on Red1 binding ^5^ and the proximity of *CEN1* to the fusion sites, this analysis was restricted to fusion chromosomes with *CEN4* deleted. Error bars show standard error. n.s. – not significant, t-test with Benjamini-Hochberg correction.

**Supplementary Fig. 6.**
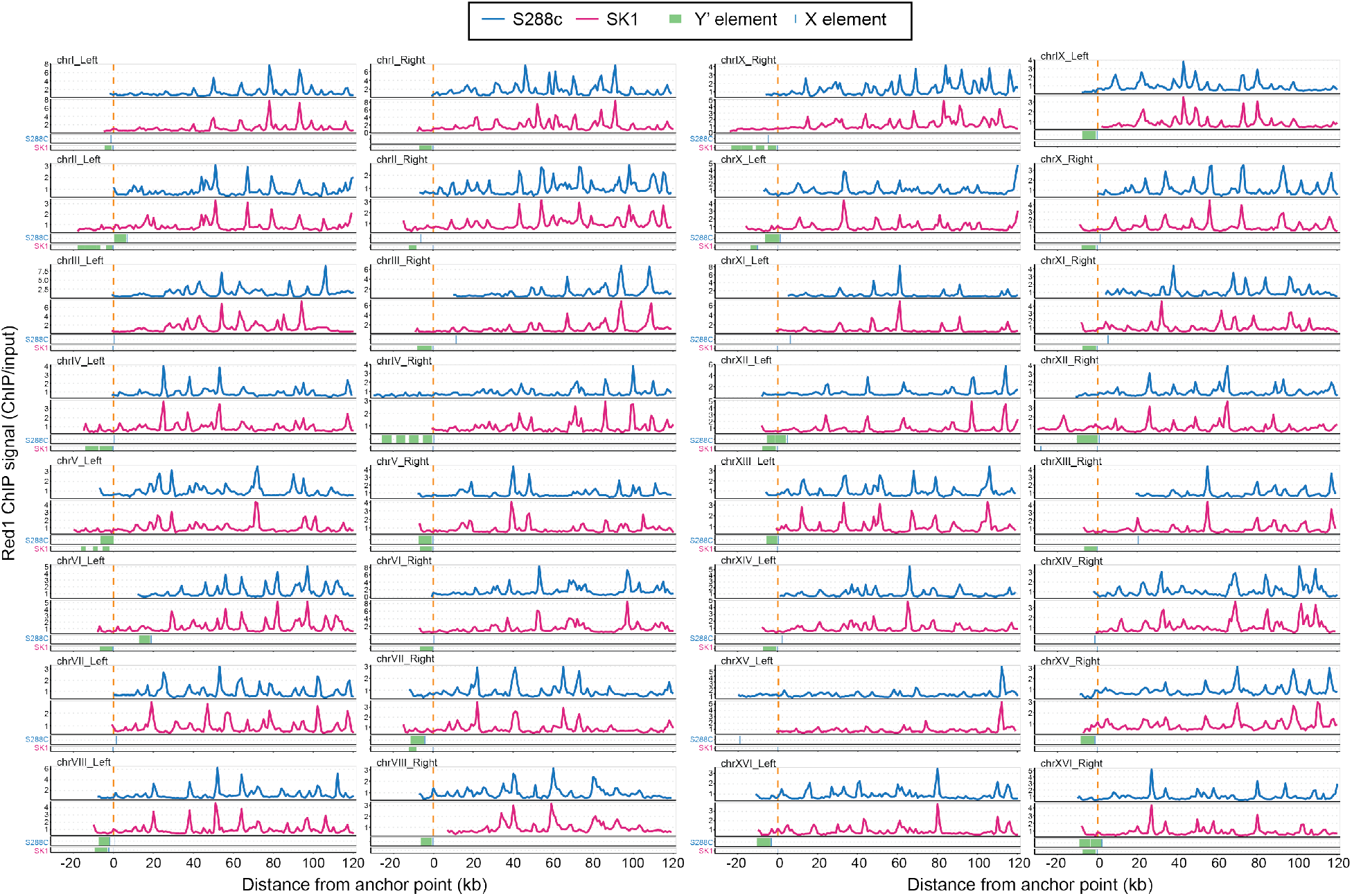
: Red1 distribution near the ends of SK1 and S288c chromosomes. Distance-from-telomere ChIP/Input signal for Red1 (T = 4h) measured in wild-type SK1/S288c hybrid strains carrying a haploid genome of SK1 and a haploid genome of S288c ^5^. The sequences between SK1 and S288c are sufficiently different that about 25% of reads can be assigned to one of the two genomes ^3^. The corresponding ends for S288c (blue) and SK1 (red) are shown juxtaposed for all 32 ends. Profiles were computationally placed in register using peak distribution and anchored on the X element annotated in SK1 as described in the Methods. Anchor points are indicated as dashed orange lines. In the tracks below the profiles, green blocks mark all annotated Y′ elements and blue tick mark all annotated X elements in S288c and SK1, respectively.

**Supplementary Fig. 7.**
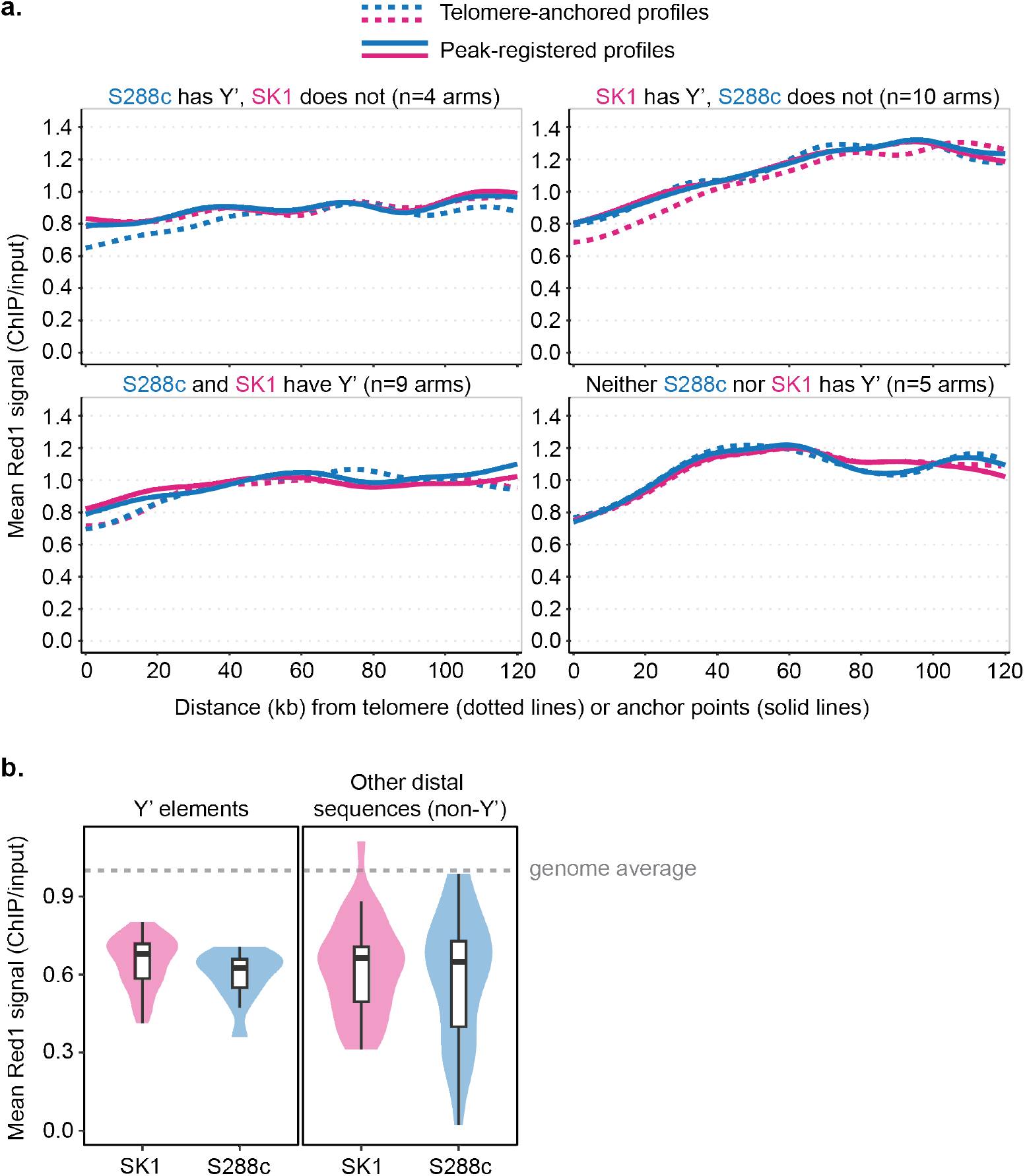
: Red1 distribution in SK1 and S288c separated by the presence of Y′ elements. **(a)** Mean distance-from-telomere ChIP/Input signal for Red1 (T = 4h) measured in wild-type SK1/S288c hybrid strains carrying a haploid genome of SK1 and a haploid genome of S288c ^5^. Panels show averages of chromosome arms depending on whether one or the other, both, or neither chromosome arm encodes Y′ elements, as labeled. The number of chromosome arms averaged in each class are indicated. In each panel, the dashed lines show mean distance-from-telomere Red1 signal anchored at the respective chromosome ends, whereas the solid lines show mean distance-from-telomere Red1 signal anchored on the SK1 X element after placing chromosome arms in register using enrichment peaks. Registered anchors are indicated as orange dashed lines in **Supplementary** Fig. 6. **(b)** Mean enrichment of Red1 on Y′ elements and on other distal sequences (i.e. sequences located telomere-proximal from the anchor points) in SK1 and S288c. Violin plots show the full distribution of means and are normalized to a genome average of 1 (gray dashed line). Embedded boxplots mark the median and interquartile range.

**Supplementary Fig. 8.**
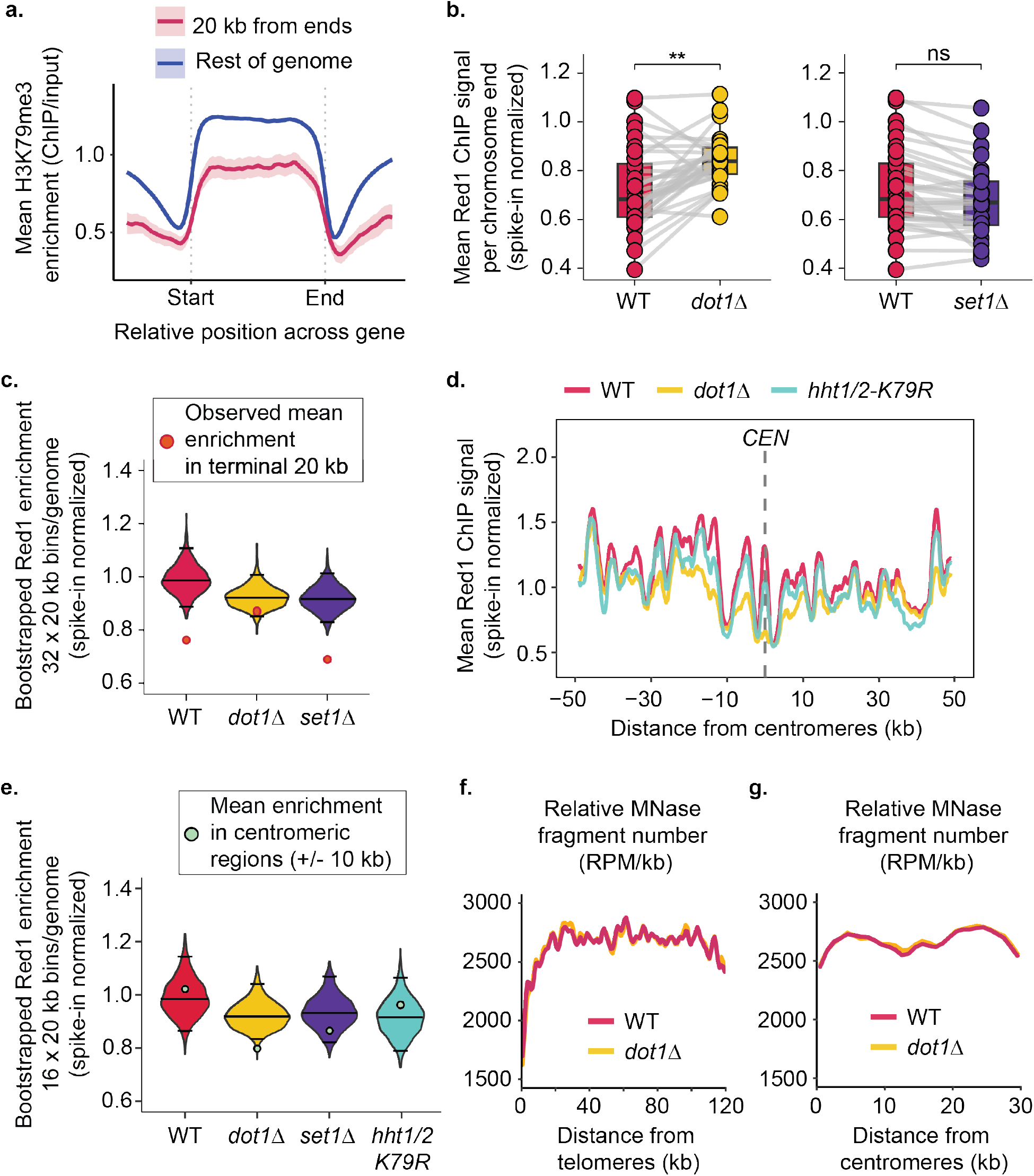
: Chromatin features and the role of histone methyltransferases at chromosome ends and centromeres. **(a)** Meta-ORF profiles of H3K79me3 (normalized to H3) in WT at early prophase I (T = 3h). Curves show mean ChIP/Input with shaded 95% CIs for genes in the terminal 20 kb (pink) versus the rest of the genome (blue; see Methods: Meta gene analyses, and meta-X and Y′ elements plots). **(b)** Mean Red1 signal per chromosome end (terminal 20 kb) in WT, *dot1Δ*, and *set1Δ* using published datasets ^2,3^. Points are individual ends. Gray lines connect matched ends across strains. Two-sided Student’s *t*-tests with BH correction; effect sizes are Cohen’s *d* (positive = higher than WT): *dot1Δ* vs WT (*p* = 0.0020, BH = 0.0038, *d* = 0.82); *set1Δ* vs WT (*p* = 0.181, BH = 0.181, *d* = 0.34). **(c)** Genome-wide bootstrap distributions of Red1 enrichment (32 × 20-kb windows; *n* = 1,000 resamples). Black lines mark medians and two-sided 95% CIs; orange/red circles mark the observed mean in the terminal 20 kb (see Methods: Bootstrap test and violin plots). Two-sided empirical tests with BH correction; Cohen’s *d* (negative = depletion at ends): WT (*p* = 0.001, BH = 0.0015, *d* = −4.15); *dot1Δ* (*p* = 0.163, BH = 0.163, *d* = −1.34); *set1Δ* (*p* < 1×10⁻⁶, BH < 1×10⁻⁶, *d* = −4.86). **(d)** Red1 distance-from-centromere profiles (± 50 kb) in WT, *dot1Δ*, and *hht1/2-K79R* (spike-in normalized). Dashed line marks the centromere midpoint. Signals were extracted in 100-bp bins around each centromere. **(e)** Bootstrap test centered on centromeres (± 10 kb; 16 × 20-kb genome windows). Two-sided empirical tests with BH correction; Cohen’s *d* (negative = depletion at chromosome ends): WT (*p* = 0.679, BH = 0.679, *d* = 0.43); *dot1Δ (p* = 0.020, BH = 0.060, *d* = −2.44)*; set1Δ* (*p* = 0.253, BH = 0.379, *d* = −1.12); *hht1/2-K79R* (*p* = 0.520, BH = 0.679, *d* = 0.65). **(f-g)** MNase-seq fragment frequency (RPM/kb) versus distance from telomeres **(f)** or centromeres **(g)** in WT and *dot1Δ*. All values are averages of two biological replicates and were reproducible across replicates.

**Supplementary Fig. 9.**
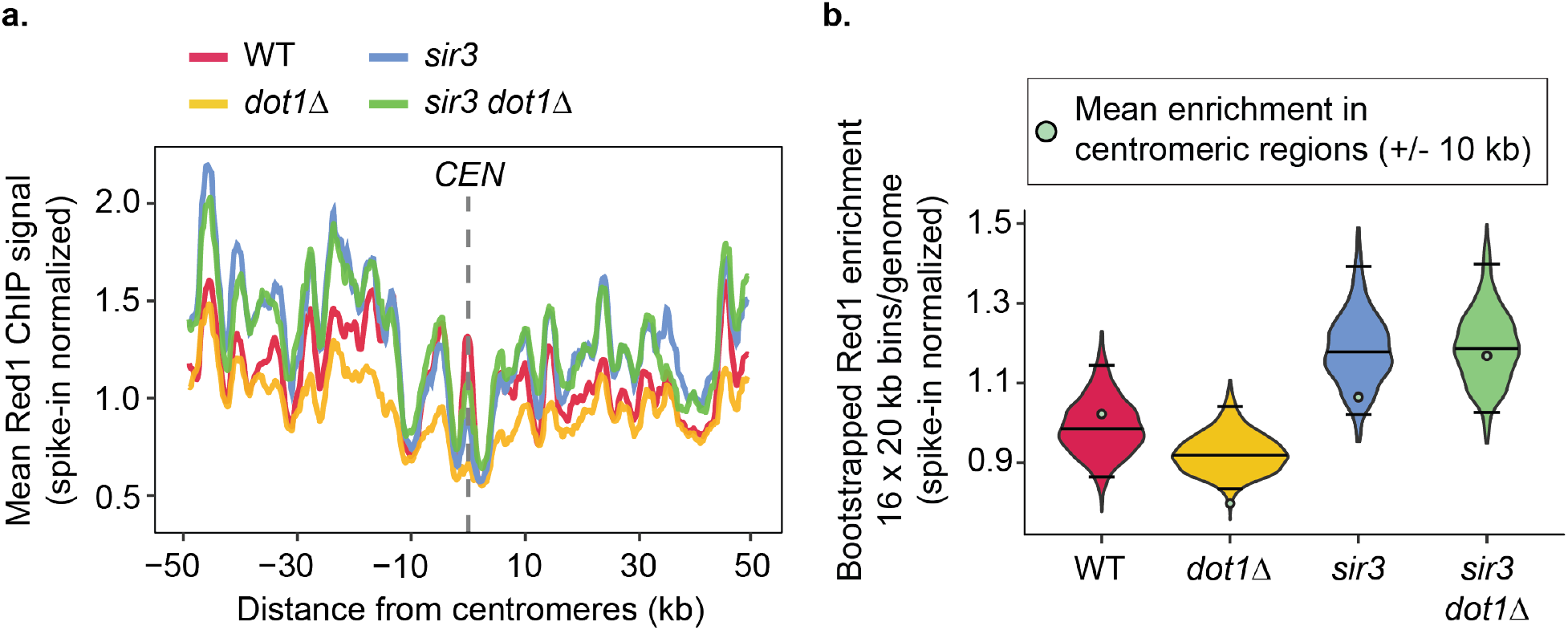
: Effects of Dot1 on axis protein deposition at centromeres depend on Sir3. **(a)** Distance-from-centromere meta-profiles (±50 kb) of Red1 ChIP in WT, *dot1Δ*, *sir3*, and *sir3 dot1Δ* (spike-in normalized). WT and *dot1Δ* are from published datasets ^2,3^. Dashed line marks the centromere midpoint. Signals were extracted in 100-bp bins around each centromere. **(b)** Genome-wide bootstrap distributions of fold-enrichment (16 × 20-kb windows; n = 1,000 resamples; see Methods: Bootstrapping plots). Black lines show medians and 95% CIs; orange/red circles mark the observed mean around the centromeres. Two-sided empirical test, effect sizes via Cohen’s d (negative = depletion at centromeres relative to the genome-wide null): WT (p = 0.679, BH = 0.797, d = +0.43); *dot1Δ* (p = 0.020, BH = 0.080, d = −2.44); *sir3* (p = 0.186, BH = 0.372, d = −1.29); *sir3 dot1Δ* (p = 0.797, BH = 0.797, d = −0.25). All values are averages of two biological replicates and were reproducible across replicates.

**Supplementary Fig. 10.**
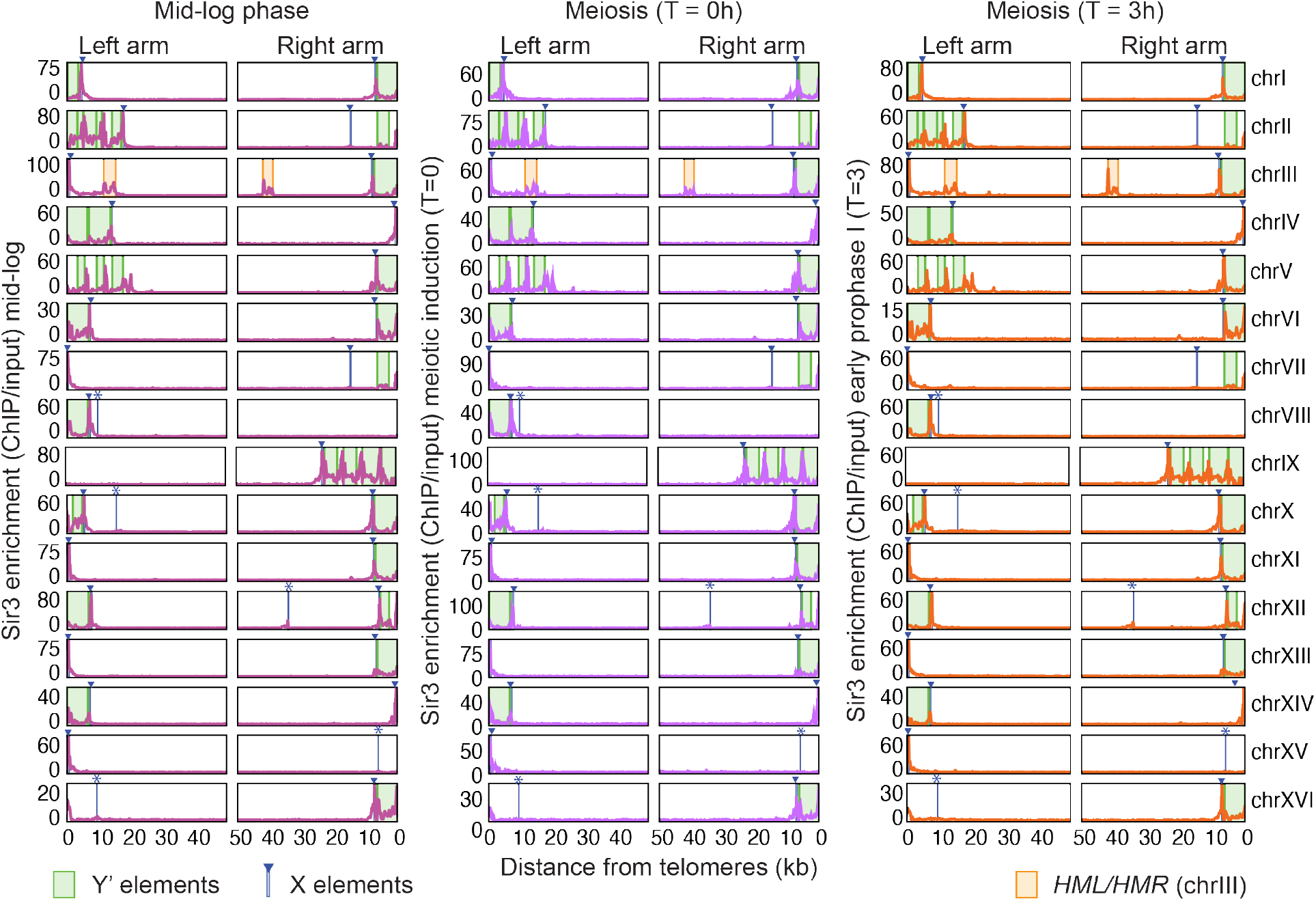
: Sir3 ChIP profiles across all chromosome ends. Distance-from-telomere Sir3 ChIP/Input signal is shown for every left and right arm (32 ends) in vegetative cells (mid-log phase), at meiotic induction (T=0), and in early prophase I (T = 3h). Green blocks mark all annotated Y′ elements and blue blocks mark X elements (fully and partially annotated copies shown: X, n = 32; Y′, n = 31). The positions of the silent mating type loci near the left (*HML*) and right (*HMR*) telomere of chrIII are also indicated. Values represent the mean of two biological replicates and were reproducible across replicates.

**Supplementary Fig. 11.**
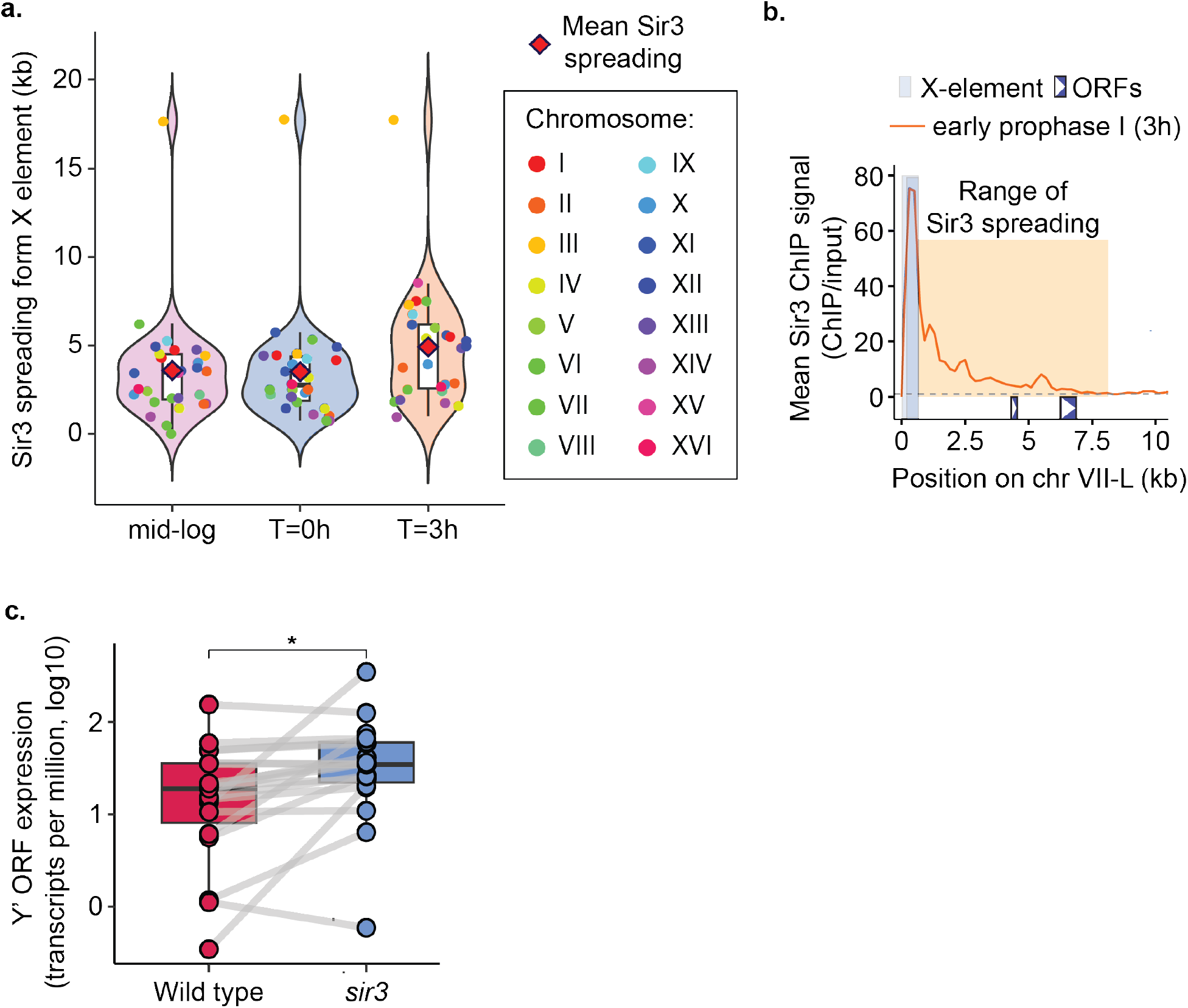
: Sir3 spreading and Y′ transcription near chromosome ends. **(a)** Sir3 spreading from the X element, quantified per chromosome end from Sir3 ChIP–seq (ChIP/Input) in early prophase I (see Methods: Quantification of Sir3 spreading from chromosome ends). The consistent outlier point indicates the spreading to *HML* on chr III-L. **(b)** Example illustrating the range of Sir3 spreading as quantified in (a). **(c)** Y′-element mRNA abundance at T = 3h after meiotic induction in WT and s*ir3* (both *ndt80Δ*). Expression is log_10_(TPM) for annotated Y′ ORFs. Points are individual Y′ ORFs and gray lines connect the same ORF across strains. Two-sided Wilcoxon rank-sum test with rank-biserial effect size (positive = higher in *sir3*) (p = 0.037, r = 0.40). Values represent the mean of two biological replicates and were reproducible across replicates.

**Supplementary Figure 12.**
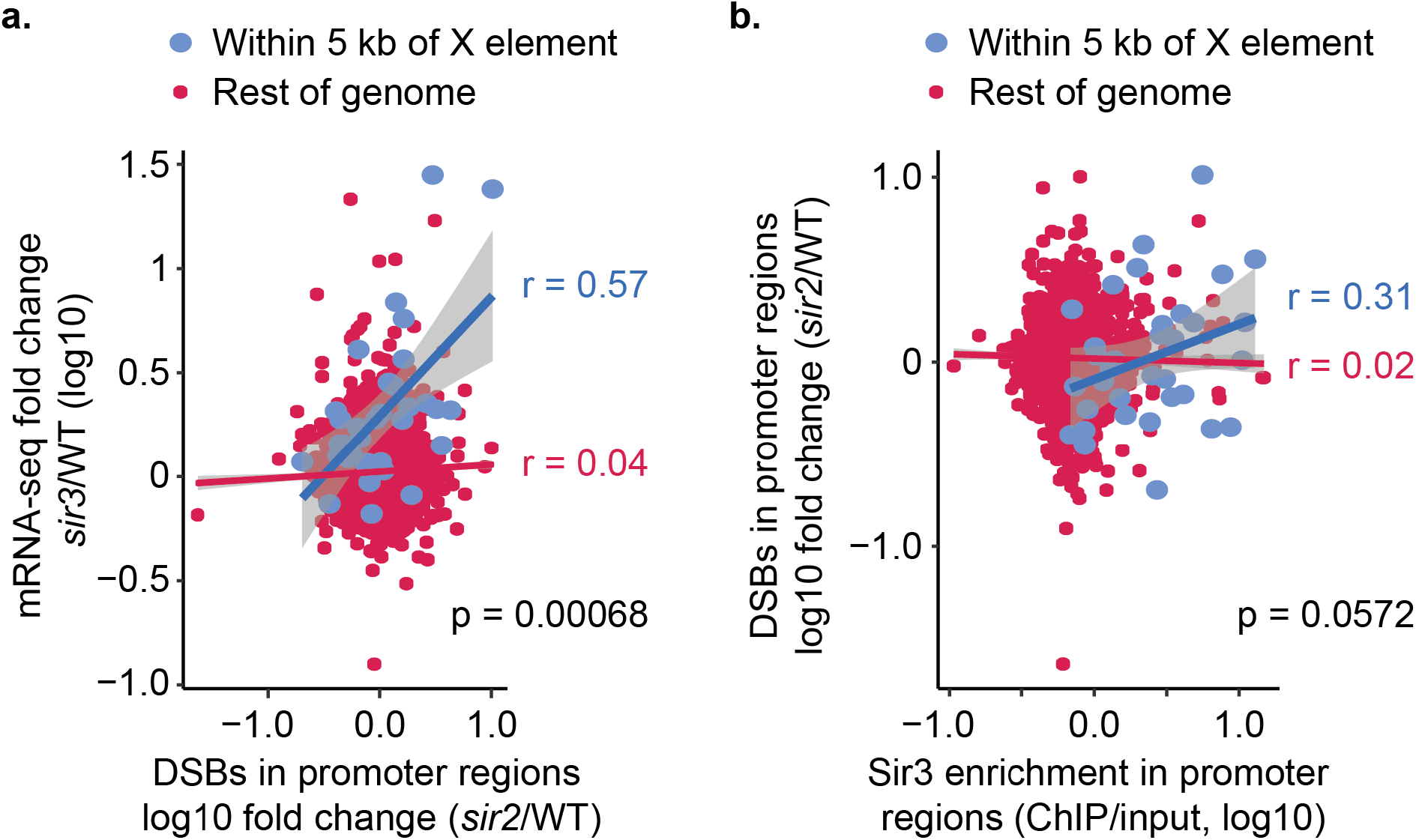
: Relationships among DSB changes, Sir3 occupancy, and transcription in promoters. **(a)** For each annotated gene, the change in Spo11-oligo signal in *sir2Δ* versus WT within the promoter (250 bp upstream of the TSS) is plotted against the change in mRNA in *sir3* versus WT (log_10_ fold change). Blue: genes within 5 kb of an X element; red: all other genes. Lines are least-squares fits with shaded 95% CIs; Pearson r values are shown. P values are based on Fisher’s r-to-z transformation and two-sided z-test. Spo11 datasets are from published work ^1,4^ (see Methods: Promoter correlation plots). **(b)** For the same promoters, DSB change (*sir2Δ*/WT, log_10_) is plotted versus Sir3 binding in WT promoters (log_10_ ChIP/Input). Color scheme and statistics as in **(a)**.

**Supplementary Figure 13.**
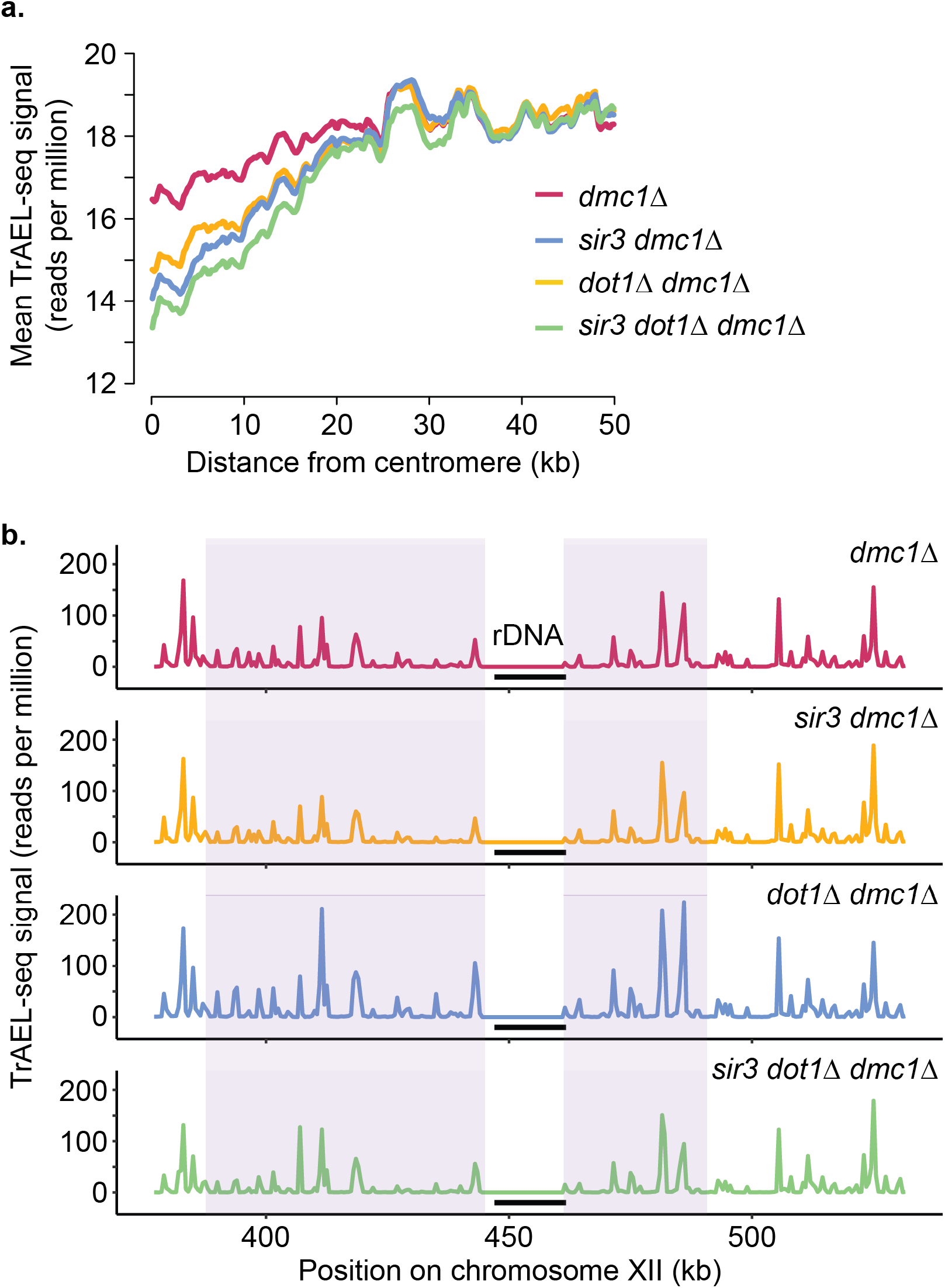
: Distinct and combined roles of Dot1 and Sir3 in shaping DSB landscapes around centromeres and the rDNA locus. (a) TrAEL-seq meta-plots showing DSB signal as a function of distance from centromeres plotted for *dmc1Δ, sir3 dmc1Δ, dot1Δ dmc1Δ,* and *sir3 dot1Δ dmc1Δ* strains. Mean TrAEL-seq signal (reads per million) plotted for the four strains within ± 50 kb of centromeres. (b) TrAEL-seq signal versus distance from the rDNA array on chr XII; the black bar denotes the rDNA repeat region. The same four strains are shown. Purple shading marks intervals with elevated DSB signal in *dot1Δ.* Values are means of two independent biological replicates and were reproducible between replicates.

## Supplementary Tables

**Supplementary Table 1.** : SK1 yeast strains utilized in this study

| Strain number | Genotype |
| --- | --- |
| AH7797 | *MATa, ho::LYS2, lys2, ura3, leu2::hisG, his3::hisG, trp1::hisG*<br>*MATalpha, ho::LYS2, lys2, URA3, LEU2, HIS3, TRP1* |
| AH5217 | *MATa, ho::LYS2, lys2, leu2::hisG, trp1::hisG, ura3, ARG4, his3::hisG, his4B::LEU2, dmc1Δ::ARG4*<br>*MATalpha, ho::LYS2, lys2, leu2::hisG, TRP1, ura3, arg4-Bgl2, his4B::LEU2, dmc1Δ::ARG4* |
| AH3083 | *MATa, ho::LYS2, lys2, leu2::hisG, his4X::LEU2-URA3, ura3, his3::hisG, trp1::hisG, arg4-Bgl2, dmc1Δ::ARG4, dot1Δ::TRP1*<br>*MATalpha, ho::LYS2, lys2, leu2::hisG, his4B::LEU2, HIS3, trp1::hisG, ura3, arg4-nsp, dmc1Δ::ARG4, dot1Δ::TRP1* |
| AH2908 | *MATa, ho::LYS2, lys2, leu2::hisG, his4B::LEU2, ura3, arg4-Bgl2, dmc1Δ::ARG4, sir3::HIS3*<br>*MATalpha, ho::LYS2, lys2, leu2::hisG, his4B::LEU2, ura3, arg4-Bgl2, dmc1Δ::ARG4, sir3::HIS3* |
| AH13037 | *MATalpha, ho::LYS2, leu2::hisG, ura3, his4B::LEU2, trp1::hisG, dot1Δ::TRP1, arg4-Bgl2, dmc1Δ::ARG4, sir3::HIS3*<br>*MATa, ho::LYS2, leu2::hisG, ura3, trp1::hisG, dot1Δ::TRP1, arg4-Bgl2, dmc1Δ::ARG4, sir3::HIS3* |
| AH8967 | *MATa, ho::LYS2, lys2, ura3, LEU2, his3::hisG, trp1::hisG, sir3::HIS3, ndt80Δ::TRP1* |
|  | *MATalpha, ho::LYS2, lys2, ura3::Ylplac211::URA3, LEU2, his3::hisG, trp1::hisG, sir3::HIS3, ndt80Δ::TRP1* |
| AH6179 | *MATa, ho::LYS2, lys2, HIS, URA, LEU, TRP, ndt80Δ::TRP1*<br>*MATalpha, ho::LYS2, lys2, leu2::hisG, his4X, ura3, trp1::hisG, ndt80Δ::TRP1* |
| AH13257 | *MATa, ho::LYS2, leu2::hisG, trp1::hisG, dot1Δ::TRP1, sir3::HIS3*<br>*MATalpha, ho::LYS2, leu2::hisG, trp1::hisG, dot1Δ::TRP1, sir3::HIS3* |
| AH2861 | *MATalpha, ho::LYS2, lys2, ura3, leu2::hisG, his3::hisG, trp1::hisG, sir3::HIS3*<br>*MATa, ho::LYS2, lys2, ura3, leu2::hisG, his3::hisG, trp1::hisG, sir3::HIS3* |
| AH8104 | *MATa, lys2, ho::LYS2, trp1::hisG, his3::hisG, LEU2, URA3, dot1Δ::TRP1*<br>*MATalpha, lys2, ho::LYS2, trp1::hisG, HIS3, leu2::hisG, ura3, dot1Δ::TRP1* |
| AH13299 | *MATalpha, ho::LYS2, lys2, ura3, leu2::hisG, his4X-LEU2(BamHI)-URA3, arg4-nsp, hht1-K79R, hht2-K79R*<br>*MATa, ho::LYS2, lys2, ura3, leu2::hisG, his4B-LEU2(MluI), arg4-bgl*<br>*hht1-K79R, hht2-K79R* |

**Supplementary Table 2.** ChIP-seq datasets utilized in this study.

| Sample | Time | Antibody | Type | GEO accession numbers | References |
| --- | --- | --- | --- | --- | --- |
| Wild type | 3 hr in meiosis | Anti-Red1 | ChIP-seq | GSE69232, GSE115092 | <sup>2,3</sup> |
| Wild type | 3 hr in meiosis | Anti-Hop1 | ChIP-seq | GSE105111 | <sup>1</sup> |
| Rec8::3HA | 3 hr in<br>meiosis | Anti-HA | ChIP-seq | GSE69232 | <sup>2</sup> |
| SK1 WT / S288c | 3 hr in<br>meiosis | Anti-Red1 | ChIP-seq | GSE114731 | <sup>5</sup> |
| S288c chrIV-chrI,<br>*cen1Δ* / SK1<br>chrIV-chrI, *cen1Δ* | 3 hr in<br>meiosis | Anti-Red1 | ChIP-seq | GSE114731 | <sup>5</sup> |
| S288c chrIV-chrI,<br>*cen4Δ* / SK1 chrIV-chrI, *cen4Δ* | 3 hr in<br>meiosis | Anti-Red1 | ChIP-seq | GSE114731 | <sup>5</sup> |
| *rec8Δ* | 3 hr in<br>meiosis | Anti-Red1 | ChIP-seq +<br>SNPChIP-seq | GSE156040 | <sup>6</sup> |
| *hop1-phd* | 3 hr in<br>meiosis | Anti-Red1 | ChIP-seq +<br>SNPChIP-seq | GSE156040 | <sup>6</sup> |
| *hop1-phd rec8Δ* | 3 hr in<br>meiosis | Anti-Red1 | ChIP-seq +<br>SNPChIP-seq | GSE156040 | <sup>6</sup> |
| Wild type | 4 hr in<br>meiosis | Anti-H4K44ac | ChIP-seq | GSE59005 | <sup>7</sup> |
| Wild type | 4 hr in<br>meiosis | Anti-H3K56ac | ChIP-seq | GSE59005 | <sup>7</sup> |
| Wild type | 4 hr in<br>meiosis | Anti-H4 | ChIP-seq | GSE59005 | <sup>7</sup> |
| Wild type | 4 hr in meiosis | Anti-H3 | ChIP-seq | GSE59005 | <sup>7</sup> |
| Wild type | 4 hr in meiosis | Anti-H3K4me3 | ChIP-seq | GSE67907 | <sup>8</sup> |
| Wild type | 3 hr in meiosis | Anti-H3K79me3 | ChIP-seq | GSE287129 | This study |
| *dot1Δ* | 3 hr in meiosis | Anti-Red1 | ChIP-seq + SNPChIP-seq | GSE115092 | <sup>2,3</sup> |
| *set1Δ* | 3 hr in meiosis | Anti-Red1 | ChIP-seq + SNPChIP-seq | GSE115092 | <sup>2,3</sup> |
| Wild type | Mid-log phase | Anti-Sir3 | ChIP-seq | GSE287129 | This study |
| Wild type | 0 hr in meiosis (post-induction) | Anti-Sir3 | ChIP-seq | GSE287129 | This study |
| Wild type | 3 hr in meiosis | Anti-Sir3 | ChIP-seq | GSE287129 | This study |
| *sir3* | 3 hr in meiosis | Anti-Red1 | ChIP-seq + SNPChIP-seq | GSE287129 | This study |
| *sir3 dot1Δ* | 3 hr in meiosis | Anti-Red1 | ChIP-seq + SNPChIP-seq | GSE287129 | This study |
| *hht1-K79R, hht2-K79R* | 3 hr in meiosis | Anti-Red1 | ChIP-seq + SNPChIP-seq | GSE287129 | This study |

**Supplementary Table 3.**
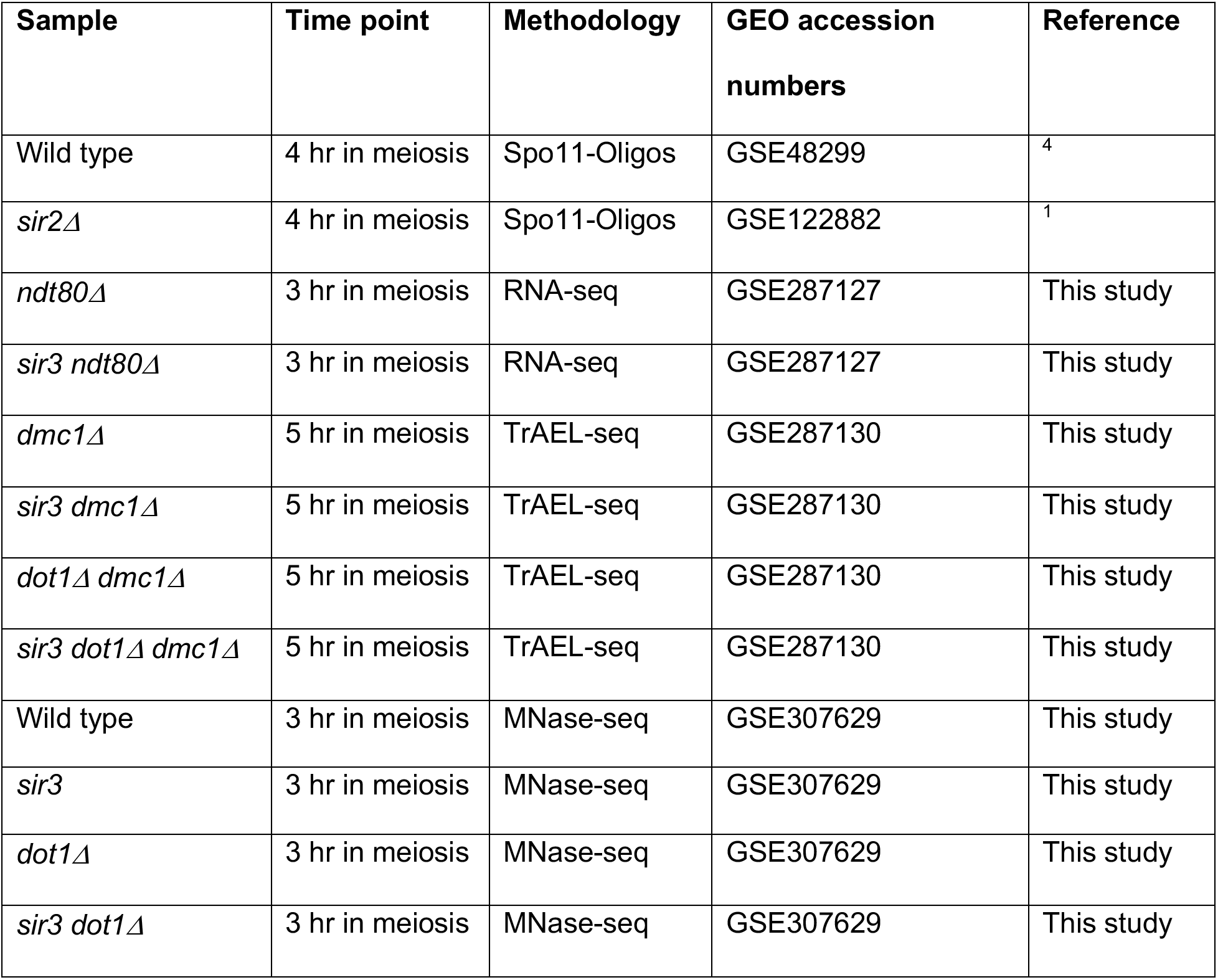
Spo11-Oligos, RNA-seq, TrAEL-seq and MNase-seq datasets utilized in this study.

| Sample | Time point | Methodology | GEO accession numbers | Reference |
| --- | --- | --- | --- | --- |
| Wild type | 4 hr in meiosis | Spo11-Oligos | GSE48299 | <sup>4</sup> |
| *sir2Δ* | 4 hr in meiosis | Spo11-Oligos | GSE122882 | <sup>1</sup> |
| *ndt80Δ* | 3 hr in meiosis | RNA-seq | GSE287127 | This study |
| *sir3 ndt80Δ* | 3 hr in meiosis | RNA-seq | GSE287127 | This study |
| *dmc1Δ* | 5 hr in meiosis | TrAEL-seq | GSE287130 | This study |
| *sir3 dmc1Δ* | 5 hr in meiosis | TrAEL-seq | GSE287130 | This study |
| *dot1Δ dmc1Δ* | 5 hr in meiosis | TrAEL-seq | GSE287130 | This study |
| *sir3 dot1Δ dmc1Δ* | 5 hr in meiosis | TrAEL-seq | GSE287130 | This study |
| Wild type | 3 hr in meiosis | MNase-seq | GSE307629 | This study |
| *sir3* | 3 hr in meiosis | MNase-seq | GSE307629 | This study |
| *dot1Δ* | 3 hr in meiosis | MNase-seq | GSE307629 | This study |
| *sir3 dot1Δ* | 3 hr in meiosis | MNase-seq | GSE307629 | This study |

